# Npl3 functions in mRNP assembly by recruitment of mRNP components to the transcription site and their transfer onto the mRNA

**DOI:** 10.1101/2022.07.22.501171

**Authors:** Philipp Keil, Alexander Wulf, Nitin Kachariya, Samira Reuscher, Kristin Hühn, Ivan Silbern, Janine Altmüller, Ralf Stehle, Kathi Zarnack, Michael Sattler, Henning Urlaub, Katja Sträßer

**Affiliations:** Institute of Biochemistry, FB08, Justus Liebig University Giessen, Heinrich-Buff-Ring 17, 35392 Giessen, Germany; Max Planck Institute for Multidisciplinary Sciences, Am Fassberg 11, 37077 Goettingen, University Medical Center Goettingen, Institute of Clinical Chemistry, Robert-Koch-Strasse 40, 37075 Göttingen, Germany; Bavarian NMR Center (BNMRZ), Department of Chemistry, Technical University of Munich, Lichtenbergstrasse 4, 85748 Garching; Institute of Structural Biology, Helmholtz Center Munich, Ingolstaedter Landstrasse 1, 85764 Neuherberg, Germany; Buchmann Institute for Molecular Life Sciences (BMLS) & Faculty of Biological Sciences, Goethe University Frankfurt, Max-von-Laue-Straße 15, 60438 Frankfurt a.M., Germany; Cologne Center for Genomics (CCG), University of Cologne, Weyertal 115b, 50931 Cologne, Germany

## Abstract

RNA-binding proteins (RBPs) control every RNA metabolic process by multiple protein-RNA and protein-protein interactions. Their roles have largely been analyzed by crude mutations, which abrogate multiple functions at once and likely impact the structural integrity of the large messenger ribonucleoprotein particle (mRNP) assemblies, these proteins often function in. Using UV-induced RNA-protein crosslinking and subsequent mass spectrometric analysis, we first identified more than 100 *in vivo* RNA crosslinks in 16 nuclear mRNP components in *S. cerevisiae*. For functional analysis, we chose Npl3, for which we determined crosslinks in its two RNA recognition motifs (RRM) and in the flexible linker region connecting the two. Using NMR and structural analyses, we show that both RRM domains and the linker uniquely contribute to RNA recognition. Interestingly, mutations in these regions cause different phenotypes, indicating distinct functions of the different RNA-binding domains of Npl3. Notably, the *npl3-Linker* mutation strongly impairs recruitment of several mRNP components to chromatin and incorporation of further mRNP components into nuclear mRNPs, establishing a function of Npl3 in nuclear mRNP assembly. Taken together, we determined the specific function of the RNA-binding activity of the nuclear mRNP component Npl3, an approach that can be applied to many RBPs in any RNA metabolic process.

## INTRODUCTION

Every RNA metabolic process entails a multitude of protein-RNA as well as protein-protein interactions. Consequently, the molecular mechanisms of these processes are difficult to study, and the molecular roles of the interactions with RNA or other proteins for a given RNA-binding protein (RBP) are known in only a few cases. In addition, many newly identified RBPs do not contain a classical RNA-binding domain (1-3). To identify their RNA-binding sites with amino acid resolution several high-throughput methods based on UV RNA-protein crosslinking coupled with mass spectrometry (MS) have been developed (4-8). The identified amino acids can then be mutated to abrogate and study the function of RNA binding of each RBP.

A prime example for an RNA metabolic process is the expression of protein-coding genes. First, the pre-mRNA is synthesized by RNA polymerase II (RNAPII) transcribing a protein-coding gene. Already co-transcriptionally, the pre-mRNA is processed, *i*.*e*. capped, spliced, cleaved and polyadenylated at its 3’ end. In addition, the mRNA is assembled into a messenger ribonucleoprotein particle (mRNP) by association with multiple RBPs (9-13). The fully processed and assembled mRNP is exported to the cytoplasm, where the mRNA is translated by the ribosomes. All these processes are mediated and coordinated by a multitude of RBPs that accompany the mRNA throughout its life cycle (14,15). Importantly, malfunction of an RBP often causes disease (11,16-19).

Nuclear mRNP assembly controls mRNA stability, nuclear mRNA export and also often regulates cytoplasmic processes such as translation, mRNA localization and degradation (9-11,20). In the model organism *S. cerevisiae*, the key RBPs involved in nuclear mRNP assembly have been identified. All of these proteins and protein complexes are conserved in higher eukaryotes, consistent with the biological importance of this process (9-11): i) The cap-binding complex (CBC) consists of the small and large subunits Cbc2 and Sto1, respectively, and promotes mRNA stability, transcription elongation, splicing, 3’ end formation and nuclear mRNA export ((21) and references therein). ii) The TREX complex couples <u>tr</u>anscription elongation to nuclear mRNA <u>ex</u>port. It is composed of the core THO complex consisting of Tho2, Hpr1, Mft1, Thp2 and Tex1, the DEAD-box RNA helicase Sub2, the mRNA export adaptor Yra1 and the two serine arginine (SR)-protein like proteins Gbp2 and Hrb1 (22,23). TREX functions in transcription elongation, 3’ end formation, mRNP assembly and nuclear mRNA export (24). iii) The <u>n</u>uclear poly(<u>A</u>)-<u>b</u>inding protein Nab2 is an SR-like protein, which functions in 3’ end processing, nuclear mRNP assembly and nuclear mRNA export ((25,26) and references therein). iv) Tho1 and its mammalian ortholog SARNP (also named CIP29) play a role in nuclear mRNP biogenesis and interact with TREX (27,28). v) The SR-like protein Npl3 functions in transcription elongation, splicing, 3’ end formation, nuclear mRNP assembly and export ((29) and references therein). Several nuclear mRNP components including Npl3, Nab2 and the TREX complex components Hpr1, Yra1, Gbp2 and Hrb1 function as mRNA export adaptors, *i*.*e*. they recruit the mRNA exporter Mex67-Mtr2 to the mRNA (11). vi) Mex67-Mtr2 transports the mRNP through the nuclear pore complex.

In summary, nuclear mRNP assembly is mediated by many RBPs, but their interplay and mechanisms of action have remained largely elusive. For example, it is unknown how each mRNP component finds its place on the mRNA, what other factors are needed for its recruitment and when this recruitment takes place. In addition, most RNP components are already recruited during transcription through protein-protein interactions, and it is not known when and how these components are transferred onto the mRNA. Furthermore, the functions of RNA binding of each RBP in nuclear mRNP assembly are unknown. A major obstacle is that these have been largely studied using crude deletion mutants that disable multiple functions and thus cause pleiotropic phenotypes due to the disruption of complex networks of many individual protein-protein and protein-RNA interactions.

Here, we used RNP^XL^ (5) combined with the purification of mRNP components to identify amino acids in RBPs that are in contact with RNA *in vivo*, and which are thus likely to mediate RNA binding. Overall, we identified more than 100 amino acid residues in mRNP components that crosslinked to RNA. We chose Npl3 to generate RNA-binding mutants as the role of its RNA-binding activity in its many different functions is unknown. Crosslinked amino acids map to the two RNA recognition motifs (RRMs), the linker region between the two RRM domains and the RG-rich domain of Npl3. Interestingly, the three generated *npl3* mutants in RRM1, the linker between the two RRM domains, or in RRM2 show different phenotypes, although each of the three mutations decreases Npl3’s ability to bind RNA. A detailed analysis of the *npl3-Linker* mutation revealed that it leads to aberrant nuclear mRNP composition and a nuclear mRNA export defect. Our results suggest that Npl3 functions to recruit the THO complex and the mRNA exporter Mex67-Mtr2 to the transcription machinery and transfers the mRNP components Sub2 and Tho1 from the transcription machinery onto the mRNA. Thus, we uncovered a novel function of Npl3 in nuclear mRNP biogenesis and the underlying molecular mechanism by the analysis of specific RNA-binding mutants. Globally, the approach presented here can be applied to any RBP in many model organisms and can thus provide novel insights into any RNA-related process.

## MATERIAL AND METHODS

### Strains, plasmids and primers

Yeast strains, plasmids and primers are listed in Supplementary Tables S9, S10 and S11, respectively. All cloning has been done by Gibson Assembly. Strains that carry a point mutation in *NPL3* have been generated by transforming a pRS315 plasmid encoding a mutated ORF + 500 bp of 5’ and 300 bp of 3’ UTR of *NPL3* into the *NPL3* shuffle strain and shuffling out the pRS316-*NPL3* by streaking cells two times over 5’-FOA plates.

### Identification of protein-RNA crosslinking sites

To purify mRNP components for the identification of RNA-binding sites a PCR amplified TAP- or FTpA-tag was genomically integrated. The PCR construct was transformed into yeast strain RS453 and verified by Western blot. For 4-thiouracil (4-tU) labeling and *in vivo* UV crosslinking, yeast cells were grown in SDC-URA supplemented with 120 µM uracil to OD_600_ = 0.8 before 4-tU was added to final concentration of 500 µM. Cells were pelleted at 7,000 g for 4 min when reaching OD_600_ ≈ 3. Cells were crosslinked by 365 nm UV light in a petri dish on a water bath with crushed ice using a Bio-Link BLX-365-UV-Crosslinker. Cells were pelleted again, suspended in 2.5 ml TAP-buffer (50 mM TRIS, pH 7.5, 1.5 mM MgCl_2_, 200 mM KCl, 0,15 % NP-40, 1 mM DTT) and flash frozen in liquid nitrogen.

Cell beads were ground using a freezer mill 6870D (SPEX SamplePrep). Lysate was cleared at 165,000 g for 1 h at 4°C. Supernatant was incubate with IgG bead-slurry (IgG Sepharose 6 Fast Flow, GE Healthcare) for 1.5 h at 4 °C. After washing, TEV-protease was added to the beads and incubated for 1 h at 16°C for elution. 100 µg of affinity-purified eluate were further processed to enrich for crosslinked protein-RNA heteroconjugates as described elsewhere (5). The enriched samples were subjected to liquid chromatography followed by mass spectrometry (LC-MS, see below). Protein-RNA crosslinks were identified by searching MS spectra with the software tool RNPxl (5,7) and NuXl (Urlaub goup unpublished) in the OpenMS environment against a database containing the FASTA sequences of the proteins Npl3, Sto1, Cbc2, Tho2, Hpr1, Mft1, Thp2, Tex1, Gbp2, Hrb1, Sub2, Yra1, Yra2, Tho1, Nab2, Sac3, Cdc31,Sus1, Sem1, Thp1, Mex67, Mtr2, Pab1, Cbf5, Mud1, Mud2, Msl5, Nam8, Prp19, Prp11, Prp42, Snu56, Xrn1, Ist3 and Yhc1 in the background of the Contaminant Repository for Affinity Purification (the CRAPome) of *S. cerevisiae* (30). In addition, MS data were searched against the entire *S. cerevisiae* database (UniProt) (Supplementary Table S3). The specific MS settings for protein-RNA crosslinking identification are described in detail in (5).

### Dot spots

Freshly grown yeast cells were picked and suspended in 1 ml water. To ensure that all spotted strains have the same number of cells the OD_600_ was measured and diluted to 0.15. A 10-fold serial dilution was made four times and 5 µl of each strain and dilution was spotted on respective media plate.

### Growth curves

An overnight culture was diluted to an OD_600_ of 0.2 and were grown for ∼2 h before the measurement of the growth curve started. To keep cells in mid-log growth phase cells were always diluted before reaching OD_600_ = 1. For incubation at 37°C a shaking water bath was used.

### RNA immunoprecipitation (RIP)

FTpA- or TAP-tagged *S. cerevisiae* strains were grown in 400 ml YDP to an OD_600_ of 0.8, harvested and stored at -80°C. Pellets were thawed on ice, resuspended in 1 ml RNA IP-buffer (25 mM TRIS HCl pH 7.5, 150 mM NaCl, 2 mM MgCl_2_, 0.2 % Triton X-100, 500 µM DTT) + protease inhibitor and lysed using the FastPrep-24 5G device (3 times for 20 s at 6 m/s). The lysate was cleared by centrifugation for 5 min at 1,500 g and 10 min at 16,000 g at 4°C. 900 µl of the cleared lysate (input) was incubated with 660 units DNase I for 30 min on ice. 40 µl IgG-coupled Dynabeads M-280 were added and incubated for 3 h at 4°C on a turning wheel. The beads were washed 8 times with RNA-IP-buffer. For RNA extraction 1 ml TRIzol reagent was added to the beads. For protein analysis, an acetone precipitation of the phenol phase and interphase was performed. The RNA of the input and IP samples were reverse transcribed and subsequently analyzed by quantitative PCR on an Applied Biosystems StepOnePlus cycler using Applied Biosystems Power SYBRGreen PCR Master Mix. As control the RIP experiment was also performed with an untagged strain (nc). PCR efficiencies (E) were determined with standard curves. Protein enrichment over the untagged strain was calculated according to 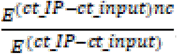. Mean values were calculated of at least three biological replicates.

### Fluorescence *in situ* hybridization with oligo d(T)

*In situ* hybridization against poly(A)+ RNA was done according to (31). Briefly, cells were grown at 30°C in YPD medium to mid-log phase before shifted to 37°C for 1 h. An aliquot of 10 ml cell culture was removed before temperature shift. Cells were immediately fixed with 4% formaldehyde, washed after fixation, and spheroplasted with 100T zymolyase. After adhering spheroplasts to poly-lysine-coated slide, the cells were prehybridized for 1 h at 37°C in prehybridization buffer (50% formamide, 10% dextran sulphate, 125 µg/ml of *E. coli* tRNA, 500 µg/ml hering sperm DNA, 4x SSC, 0.02% polyvinyl pyrrolidone, 0.02% BSA, 0.02% Ficoll-40) in a humid chamber. To hybridize with oligo d(T)50-Cy3 0.75 µl of 1 pmol/µl probe was added and incubated at 37°C overnight in humid chamber. After hybridization, cells were washed, mounted with ROTI®Mount FluorCare DAPI, and covered with a coverslip. Cells were inspected either with an Axio observer fluorescence microscope (Zeiss) connected to a CCD camera.

### Identification and quantification of affinity-purified proteins using MS

TAP purifications of Cbc2 were essentially done as described in “Identification of protein-RNA crosslinking sites”. The TEV eluates were analyzed by MS and quantitative Western blot. For MS analysis, samples were processed identically according to a modified protocol (32) to ensure equal sample quality and comparability. Briefly, proteins contained in the eluates were subjected to SDS-PAGE analysis on a 4-12% gradient gel (NuPage, Thermo Fisher Scientific) according to the manufacturer’s specifications. Next, individual lanes of eluates were cut into 23 pieces and proteins were alkylated with 55 mM iodacetamide before digested with trypsin (12ng/µl) in-gel overnight at 37°C. Extracted peptides were dried by vacuum centrifugation and resuspended in 2% acetonitrile, 0.05% TFA before subjected to LC-MS analysis using a self-packed C18 column (Dr. Maisch GmbH, 1.9 µM pore size). All samples were analyzed on a 58-minute linear gradient, the effective peptide separation was achieved through a linear increase of acetonitrile from 8 to 36% in the mobile phase over 43 min. Eluted peptides were analyzed by QExactive HF or Fusion Tribrid mass spectrometer (Thermo Fisher Scientific). MS1 spectra were acquired using Orbitrap mass analyzer at the resolution of 60,000 (QExactive HF) or 120,000 (Fusion Tribrid). MS2 spectra were acquired in data dependent mode using Orbitrap at 15,000 resolution (QExactive HF) or IonTrap in the rapid acquisition mode (Fusion Tribrid). Raw spectra were queried against reviewed canonical sequences of *S. cerevisiae* (retrieved from UniProt in August 2020, 6,049 entries), supplemented with a *npl3-P196D/A197D* (*npl3-Linker*) modified sequence. Processing of Cbc2-TAP samples together with samples derived from the Npl3-TAP purification with and without RNase A treatment (not shown) was accomplished by MaxQuant version 1.6.2.10 (33) using default settings, except: i) maximum peptide mass was increased to 6000 Da; ii) enabled “Match between runs” and calculation of iBAQ values (34). Only intensities of proteins with at least two quantified razor/unique peptides in one of the affinity purification experiments were considered for further analysis. Missing iBAQ values were imputed by random sampling from a normal distribution with a center at 20% percentile and 0.5 standard deviation of iBAQ intensity in the sample. Raw iBAQ intensities were log-transformed and median-normalized. Proteins enriched in mock-control as compared to the whole lysates were not considered as candidate binders. iBAQ intensities in affinity-purified samples were normalized to the abundances of the bait protein (Cbc2). Differences in the protein abundance between *npl3* mutant and *NPL3* wild-type (wt) were tested using limma. Empirical-Bayes modified p-values were further subjected to multiple testing correction following (35).

### Quantitative Western blots

To quantify the total amount of a protein, cells were grown to mid-log phase and lysed by FastPrep-24 5G device. For quantification equal amounts of total protein were separated by SDS-PAGE, and proteins were detected with the corresponding primary antibody, a horse radish peroxidase-coupled secondary antibody and CheLuminate-HRP ECL solution (Applichem). Chemiluminescence signals were imaged using a ChemoCam Imager (Intas) and quantified using FIJI. Mex67 and Hpr1 were HA-tagged and detected with an anti-HA antibody fused with HRP (R&D Systems). The antibody directed against Sto1 / Cbp80 is a kind gift of D. Görlich (36).

### Chromatin immunoprecipitation

Chromatin immunoprecipitation (ChIP) experiments in *S. cerevisiae* were performed according to (37) with some modifications. Briefly, 100 ml yeast culture in mid-log phase were crosslinked with 1% formaldehyde for 20 min. The reaction was stopped by adding glycine. Cells were lysed using a FastPrep-24 5G device twice for 45 s (6.5 m/s setting) with 2 min break on ice in between. The lysate was sonicated using a Bioruptor UCD-200 (Diagenode) for 3 × 15 min (30 s ON/30 s OFF) at ‘HIGH’ power setting with intermittent cooling for 5 min resulting in chromatin fragments of ∼250 bp. TAP- or FTpA-tagged proteins were immunoprecipitated with IgG-coupled Dynabeads (tosylactivated M280, Thermo Scientific) for 2.5 h at 20°C. For ChIP experiments of RNAPII, the monoclonal antibody 8WG16 (Biolegend) was added for 1.5 h at 20°C followed by 1 h incubation with Protein G Dynabeads. To reverse crosslink samples were incubated for 2 h at 37°C followed by overnight step at 65°C. To purify the DNA the PCR NucleoSpin® Gel and PCR Clean-up-kit (Macherey-Nagel) was used according to manufacturers’ manual except that the elution was done in 140 µl 1xTE. A non-transcribed region (NTR1, 174131–174200 on chr. V) served as negative control. The occupancy of each protein was calculated as its enrichment at the respective gene relative to NTR1.

### Protein expression and purification

His_6_-tagged RRM1-RRM2 tandem domains (residues 120-280) of wt Npl3 (named Npl3^120-280^) and the three mutants npl3-RRM1, npl3-Linker and npl3-RRM2 (named npl3^120-280^-RRM1, npl3^120-280^-Linker and npl3^120-280^-RRM2) were expressed in *E. coli BL21* (DE3) in minimal M9 media supplemented with ^15^N-labeled NH_4_Cl and/or ^13^C-labeled glucose as sole nitrogen and carbon source, respectively. Protein expression was induced at an OD_600_ of 0.8 with 0.5 mM isopropyl β-d-1-thiogalactopyranoside and cells grown at 22°C overnight. Cells were harvested, stored at -20°C and lysed by sonication. Proteins were purified by standard affinity chromatography with Ni-NTA sepharose. The N-terminal His-tag was removed by cleavage with TEV protease. Further purification was done using ion exchange and size exclusion chromatography. Purified samples were exchanged with NMR buffer containing 20 mM sodium phosphate, pH 6.4, 50 mM NaCl, 1 mM DTT. 10 % D_2_O was added to lock the magnetic field.

Backbone chemical shift assignments for Npl3^120-280^ were obtained from the BMRB (ID 7382 and 7383), while missing assignments for tandem RRM domains of Npl3^120-280^ and npl3^120-280^-Linker were obtained using 3D triple resonance experiments: HNCACB, HNcoCACB, HNCO, HNcaCO and HcccoNH (38). For RNA-binding studies, ^1^H-^15^N heteronuclear single quantum coherence (HSQC) and HN^ε^-selective heteronuclear in-phase single quantum coherence (HISQC) spectra (39) were recorded in a titration series using 50 μM of protein with increasing concentration of single-stranded DNA and RNA (synthetic DNA and RNA purchased from eurofins Genomics and Dharmacon, USA, or Biolegio BV, Germany, respectively). Chemical shift perturbations (CSPs, Δδ) were calculated as: Δδ =[(Δδ^1^H)^2^ + (Δδ ^15^N)^2^/25]^1/2^.

Dissociation constants (*K*_D_) were derived from NMR titrations by fitting to the following equation: Δδ_obs_ = Δδ_max_ {([P]_t_ + [L]_t_ + *K*_*D*_) - [ ([P]_t_ + [L]_t_ + *K*_*D*_)^2^ - 4[P]_t_ [L]_t_ ]^1/2^} / 2[P]_t_, where, Δδ_obs_ is the observed chemical shift difference relative to the free state, Δδ_max_ is the maximum shift change in saturation, [P]_t_ and [L]_t_ are the total protein and ligand concentrations, respectively, and *K*_D_ is the dissociation constant (40). {^1^H}-^15^N heteronuclear NOE (hetNOE) was determined from the ratio of signal intensities with and without saturation (41) in HSQC-based experiments with 3 s interscan delay. HetNOE for the protein alone using 200 to 300 µM and the protein-RNA complex were recorded with 90 µM protein and a 3-fold molar access of RNA. A series of ^1^H,^15^N HSQC experiments were recorded over 4 h to monitor amide hydrogen-deuterium exchange.

NMR measurements were carried out with NMR samples in a Shigemi tube (Shigemi Inc, Japan) at 25°C on Bruker spectrometers operation at 500, 600, 800, 900 or 950 MHz proton Larmor frequency equipped with room temperature or cryogenic probes. NMR spectra were processed with a shifted sine-bell window function and zero-filling before Fourier transformation using Bruker Topspin 3.5pl6 and NMRPipe (42). Proton chemical shifts were referenced against sodium 2,2-dimethyl-2-silapentane-5-sulfonate (DSS). All spectra were analyzed with the CCPNMR analysis v2.5 software package (43).

For Npl3^120-280, 15^N relaxation data were measured as described (41). For ^15^N, *R*_2_ pseudo-3D version of experiments were used sampling the exponential decay with CPMG pulse trains at 16.96, 33.92, 50.88, 67.84, 101.76, 135.68, 169.60, 203.52 and 254.40 ms. The signal intensity decay was fitted to an exponential decay. For longitudinal ^15^N relaxation rates, *R*_*1*_ were measured by sampling exponential decay function of delays with 60, 100, 160, 200, 280, 400, 600, 800, 1200 and 1800 ms. From the ratio of *R*_*2*_ and *R*_*1*_ rates, residue-specific rotational correlation times (τ_c_) were determined as described (44).

### Paramagnetic relaxation enhancement (PRE) experiments

Single cysteine point mutations were introduced in Npl3^120-280^ (D135C, E176C, N185C and D236C) and the native Cys211 was mutated to serine. Mutant proteins were purified as described above. Before spin labelling, samples were reduced with 5 mM DTT and dialyzed against buffer containing 50 mM Tris, pH 8.0, 50 mM NaCl. A 3-(2-lodoacetanido)-PROXYL (IPSL) stock was prepared freshly in DMSO and incubated with protein at 10-fold excess for 24 h. Excess spin label was removed by using a desalting column (PD-10, GE Heath Care) and buffer exchanged with 3 kDa cut-off Amicon filter with final NMR buffer. All steps were carried out in the dark. Labelling efficiency of proteins was confirmed by native electron spray ionization mass spectrometry. Samples were reduced by adding 10-fold excess of ascorbic acid followed by incubation for 1 h to ensure complete reduction. ^1^H paramagnetic relaxation enhancements were recorded and analyzed as described (45). HSQC and HISQC spectra were recorded with the oxidized and reduced form with a 3 s inter-scan delay for 7 h. PREs with the RNA bound form were recorded for spin labels at residue 185 and 236 with 3-fold excess of the “CN--GG” RNA oligo.

### Structure calculations

Structure calculations were carried out as described previously (45,46). In brief, structural coordinates of the individual RRM domains were taken from the Protein Data Bank (PDB) accession codes 2OSQ and 2OSR for RRM1 and RRM2, respectively. N- and C-terminal flexible linkers were removed, and only the rigid core of domains were kept for refinement. Cysteine coordinates were replaced at position of 135, 176, 185 and 236 in the PDB file, and coordinates of spin label IPSL moieties were attached to the cysteine side chain. A short molecular dynamic (MD) simulation was performed to randomize the N-terminal, C-terminal and linker between two RRM domains, and these coordinates were used as a starting template for structure calculation. TALOS-N (47) was used to generate backbone torsion angle restraints based on secondary chemical shifts. PRE-based experimental distance restraints were derived for individual dataset of 135, 176, 185 and 236 spin. Combining torsion angle and distance restraints, structural refinement was carried out using CNS 1.2. 100 randomized models were generated and analyzed. Quality factor (Q-factor) was derived by comparing the back calculated PRE curves (from the generated models) and the experimental PRE as described (45). An ensemble of the 15 lowest energy structures was selected and further analyzed. For solvent refinement of the final structures, spin label moieties were removed and native residues were replaced in the final ensemble PRE model. The position of the rest of the atoms was fixed and a short energy minimize simulation was performed. Final refinement in explicit water was performed using Aria 1.2/CNS 1.2 (48). The backbone coordinate precision of the final ensemble of structures corresponds to an RMSD of 0.93 Å. The structure is in agreement with the experimental data with an average PRE Q-factor of 0.1. The final structure was also validated with experimental SAXS data, with a χ^2^= 2.7. Backbone structural quality of the final ensemble of structures was checked by Ramachandran plot using Procheck_NMR 3.5 (49), showing 82.8% residues in allowed, 14.8% additionally allowed, 1.6% generously allowed and 0.8% in dis-allowed regions of the Ramachandran plot (Supplementary Table S6). The protein structures were visualized using PyMol by Schrodinger tool (http://pymol.org/2/).

To derive a structural model of the protein-RNA complex, we measured the PRE experiments as above with spin labels attached to 185C (on RRM1) and 236C (on RRM2) position in presence of 3-fold excess of “CN--GG” RNA. Protein-RNA distance restraints were obtained from chemical shift perturbations seen in NMR titration and based on the homologous structure (PDB: 2M8D and 5DDR). Aria/CNS1.2 was used to generate a pool of 400 models, which were then scored against experimental SAXS data (50). To do this, theoretical SAXS curves were generated using CRYSOL from the ATSAS software package 3.0.0 (51) and compared with the experimentally measured SAXS data. The final structural model of the protein-RNA complex shows a χ^2^ of 1.9 with the experimental SAXS data.

### Small angle X-ray scattering (SAXS) measurements

SAXS measurements were performed in-house on a Rigaku BIOSAXS1000 instrument mounted to a Rigaku HF007 microfocus rotational anode with a copper target (40 kV, 30 mA). Transmissions were measured with a photodiode beam stop. Calibration was done with a silver behenate sample (Alpha Aeser). Samples were measured in 12900 second frames to check beam damage. Samples were dialyzed with NMR buffer before measurement and protein-RNA complex with “CN--GG” RNA was prepared using size exclusion chromatography. To eliminate a concentration-dependent effect, a different concentration range from 2 to 8 mg/ml was measured for each dataset at 4°C. Buffer samples were measured multiple times in-between each run and applied for buffer subtraction by using SAXSLab software (v3.02). Pair-distance distribution function, *P*(r), and double logarithmic plots were calculated using the ATSAS software package 3.0.0 (52). A theoretical SAXS curves from structures were generated by CRYSOL (ATSAS software package).

### Isothermal titration calorimetry (ITC)

ITC experiments were performed with a MicroCal PEAQ-ITC device (Malvern, UK). All protein samples were dialyzed against NMR buffer (20 mM sodium phosphate, pH 6.4, 50 mM NaCl, 1 mM TCEP). The ITC cell was filled with a 15 µM concentration of RNA oligo and protein was added from a syringe. Titrations were performed with 39 points of 1 μl injections with a 150 s interval at 25°C. All measurements were conducted in duplicates and analyzed by the Malvern’s MicroCal PEAQ-ITC analysis software (v1.0.0.1259). Binding curves were fitted to one-site binding mode and thermodynamic parameters were extracted.

### Transcriptome wide analysis of differential gene expression and splicing

Total RNA was extracted with Trizol reagent (Invitrogen) according to manufactures protocol. RNA was precipitated by adding 0.5 ml 2-propanol and 2 µL glycogen (5 mg/ml) and dissolved in DEPC treated RNase-free water. For sequencing the purity and integrity of the RNA was analyzed by Bioanalyzer on an RNA Nano Chip (Agilent Technologies) according to manufactures’ manual.

After RNA quantification using the Qubit RNA BR Assay Kit, ERCC RNA Spike-In Mix (both Invitrogen) was added according to the manufacturer’s instructions. Libraries were prepared using the Illumina® TruSeq® mRNA stranded sample preparation Kit. After poly-A selection (using poly-T oligo-attached magnetic beads), mRNA was purified and fragmented using divalent cations under elevated temperature. The RNA fragments underwent reverse transcription using random primers. This was followed by second strand cDNA synthesis with DNA Polymerase I and RNase H. After end repair and A-tailing, indexing adapters were ligated. The products were then purified and amplified to create the final cDNA libraries. After library validation and quantification (Agilent 4200 Tape Station), equimolar amounts of library were pooled. The pool was quantified by using the Peqlab KAPA Library Quantification Kit and the Applied Biosystems 7900HT Sequence Detection System and subsequently sequenced on a NovaSeq6000 sequencer with 100 nt paired-end reads aiming at 25 million reads per sample.

### RNA-seq data analysis

For all genomic analyses, *S. cerevisiae* S288c genome and gene annotation assembly (version R64-1-1) were downloaded from the EnsemblFungi database (https://fungi.ensembl.org/index.html). After initial quality control using FastQC (https://www.bioinformatics.babraham.ac.uk/projects/fastqc/), RNA-seq reads were mapped using the splice-aware alignment tool STAR (version 2.7.2a) (53) with the following parameters: --outFilterMultimapNmax 1 --outFilterMismatchNoverLmax 0.04. For visualization, bam files were converted to bigwig files using deepTools (54), including count per million (CPM) normalization.

For differential gene expression analysis, reads were counted into exons using HTSeq-count from the HTSeq Python package (55). Comparison between wt (NPL3) and each of the three *npl3* mutants was performed with the R/Bioconductor package DESeq2 (version 1.28.1) (56) including shrinkage of logarithmic fold changes to disregard genes with low read counts. Log_2_-transformed fold changes of intron-containing and intron-less genes were compared using Wilcoxon rank-sum test.

Analysis of intron retention events was implemented using the Bioconductor package GenomicAlignments (version 1.24.0) (57). To quantify intron retention, the average number of reads overlapping the exon-intron (EI) 5’ and 3’ boundaries was counted for each intron. Similarly, reads overlapping the exon-exon (EE) junction were counted to evaluate spliced isoforms. The percentage of intron retention was calculated as

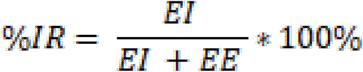

Only genes with EI + EE > 100 were considered (n = 215). Differential intron retention was analyzed between NPL3 and each of the three *npl3* mutants. For each gene, ΔIR values were calculated based as

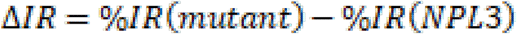

Wilcoxon rank-sum tests were performed to determine statistically significant changes in ΔIR distributions for all intron-containing genes between NPL3 and *npl3* mutants. All analyses were performed in R (version 4.0.2).

### Statistical analysis

All data are presented as average ± standard deviation (error bars) of at least three biologically independent experiments. Asterisks indicate the statistical significance (Student’s t-test; * = p-value ≤ 0.05; ** = p-value ≤ 0.01; *** = p-value ≤ 0.001).

## RESULTS

### Identification of amino acids in nuclear mRNP components that bind to RNA *in vivo*

A multitude of nuclear RBPs bind to mRNA, forming mRNPs and regulating every step of post-transcriptional gene regulation (9-11,13,58). To find out which functions are served by the RNA-binding activity of these nuclear mRNP components, we sought to disrupt specifically and individually their RNA-binding activity. First, we identified amino acids that make direct contact with RNA *in vivo* by UV crosslinking combined with MS, a method that has proved reliable in identifying RBPs and their crosslinked amino acids in several proteome-wide approaches (5,8,59-62). However, unlike in these previous studies that followed a strategy of purifying the entire (pre-m)RNA pool, we used a targeted approach by purifying specific proteins and protein complexes after UV crosslinking. To do this, we genomically TAP-tagged the following mRNP components in *S. cerevisiae*: the cap-binding complex subunit Cbc2, the TREX subunits Hpr1 and Sub2, Tho1, the SR-like proteins Nab2 and Npl3, the THSC subunit Thp1 and the mRNA exporter subunit Mtr2. In each of the TAP-tagged strains, we labeled the RNA *in vivo* with 4-thiouracil (4-tU), crosslinked RNA and proteins by UV irradiation at 365 nm and natively purified each mRNP component and any associated proteins. Purified complexes were digested with nucleases and trypsin. Non-crosslinked RNA (oligo)nucleotides were removed by C18 chromatography and crosslinked peptides were enriched by TiO2 chromatography. Amino acids in peptides crosslinked to RNA were identified by MS using the software RNP^XL^ for data analysis of the various purified mRNP complexes (Figure 1A) (5,7).

**Figure 1.**
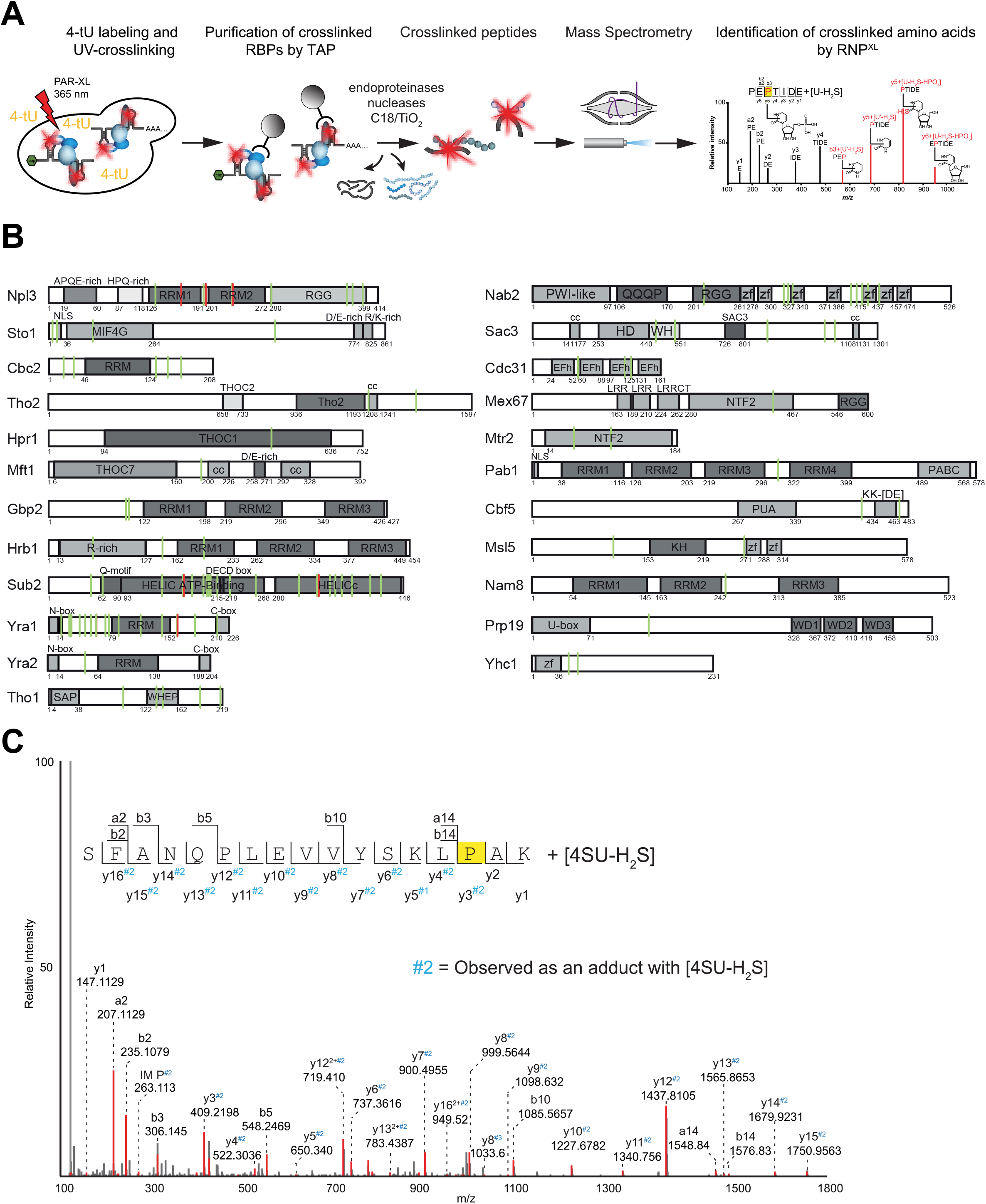
Amino acids crosslinking to RNA *in vivo* in proteins involved in nuclear mRNP assembly. (**A**) Schematic overview of the experimental workflow to identify *in vivo* protein-RNA binding sites. RNA was labeled with 4-tU and crosslinked to RNA by UV irradiation. Living yeast cells were lysed and protein complexes of interest purified by TAP. TEV eluates of UV-irradiated and control cells were digested with nucleases and endoproteinase trypsin and non-crosslinked RNA oligonucleotides were removed by C18 chromatography. Crosslinked peptide-RNA (oligo)nucleotides were enriched with TiO2 and analyzed by LC-MS. Peptides and amino acids crosslinked to RNA were identified using RNPxl (5,7). (**B**) Linear representation of the crosslinked proteins with the amino acids identified to be crosslinked (green vertical line). Each protein is depicted with its domain organization and amino acid positions. Vertical red lines indicate crosslinked amino acids in the corresponding proteins that had been previously identified by Kramer et al. (2014). Sto1, Tho2, Hpr1, Sac3 and Mex67 are not drawn to scale. Abbreviations: RRM: RNA recognition motif; RGG: arginine-glycine-rich domain; NLS: nuclear localization signal; MIF4G: middle domain of eukaryotic initiation factor 4G; THOC2: THO complex, subunit 2, N-terminal; Tho2: THO complex, subunit 2, C-terminal; THOC1: THO complex subunit 1; THOC7: THO complex subunit 7; cc: coiled coil; Q-motif: characteristic for DEAD box helicases (GFXXPXPIQ); HELIC ATP-binding: ATP-binding domain; HELICc: C-terminal helicase domain; N-box/C-box: highly conserved in REF proteins; SAP: SAF-A/B, Acinus and PIAS; WHEP: domain discovered in TrpRS (W), HisRS (H), GluProRS (EP); zf: C3H1 zinc finger; HD: superhelical domain; WH: winged helix domain; EFh; EF-hand; NTF2: nuclear transport factor 2; PABC: conserved C-terminal domain of the polyadenylate-binding protein (PABP); PUA domain: pseudouridine synthase and archaeosine transglycosylase domain; KH: K-homology domain. (**C**) MS/MS spectra of a peptide from Npl3 (amino acid positions 182 to 198) crosslinked to 4-thiouridine. Upon UV crosslinking of 4-tU a neutral loss of H_2_S is observed. The sequence of the peptide is listed and its production ions with and without the adduct mass of 4-tU minus H_2_S are indicated. The crosslinked P196 is highlighted in yellow.

In total, we identified more than 100 crosslinked peptides in 23 nuclear mRNP components and co-purified splicing factors (Figure 1B, Supplementary Tables S1 and S2). For most of these peptides we were able to pinpoint the crosslink to a single or two adjacent crosslinked amino acids. Three RBPs emerge with high numbers of crosslinked peptides as well as identified crosslinked amino acids, namely Sub2, Yra1 and Nab2 (Supplementary Table S1). Most Sub2 crosslinking sites fall into the helicase domains. Crosslinking sites in Yra1 are not only found in the RRM, but are distributed throughout the protein’s sequence. Some crosslinking sites in Nab2 are located within the RGG domain, but the majority is found within the protein region that includes the zinc-finger domains. Interestingly, we found that the wing-helix (WH) domain, which is a classical DNA-binding motif in Sac3, was crosslinked to RNA, as was the WHEP-TRS domain, a helix-turn-helix DNA-binding motif in Tho2 (Figure 1B). The identified amino acids can be specifically mutated in order to disrupt the RNA-binding activity of each mRNP component to assess its function in nuclear mRNP assembly as well as other steps of post-transcriptional gene regulation.

### mRNA-binding mutants of Npl3 cause severe growth defects

Based on its many functions in post-transcriptional gene regulation we chose Npl3, an SR-like protein found in yeast and homologous to many eukaryotic SR-like proteins, for an in-depth structural and functional analysis. Npl3 contains a canonical RRM domain (RRM1) connected by a short linker (8 amino acids) to a non-canonical RRM2 followed by an RGG domain (Figure 1B). A multiple sequence alignment of SR-like proteins across different species shows conservation of consensus ribonucleoprotein (RNP) motifs in the RRM1 domains, while the RRM2 domain of Npl3 lacks consensus RNP motifs. Instead, the RRM2 shares highly conserved motifs with other non-canonical RRM domains (Supplementary Figure S3A). We detected crosslinks between RNA and amino acids in both RRM domains, including the canonical phenylalanine (F162) in the RNP1 motif of RRM1 and F229/S230 in RRM2 (Figure 1B; Supplementary Tables S1 and S2). Additional crosslinks were found in the RGG domain. Importantly, we also identified two RNA-crosslinked amino acids in the linker region between the two RRM domains (P196 and A197) revealing an involvement of this region in RNA binding. Figure 1C shows an MS/MS spectrum of a peptide derived from Npl3 encompassing amino acid positions 182 to 198 crosslinked to 4-tU derivative, which has lost H_2_S due to the UV crosslinking. The fragment spectrum reveals a complete y-type product ion series of the peptide. Importantly, from y3 to y16 all product ions have a mass adduct corresponding to a 4-tU minus H_2_S proving that P196 is crosslinked to 4-thiouridine. To determine the function of Npl3’s RNA-binding activity and the domains involved, we mutated the amino acids identified as crosslinked to RNA (i) in each of the two RRM domains and (ii) in the linker region (Figure 2A and B and Supplementary Table S1). The selected mutants were named *npl3-RRM1, npl3-RRM2* and *npl3-Linker*, respectively (Figure 2A).

**Figure 2.**
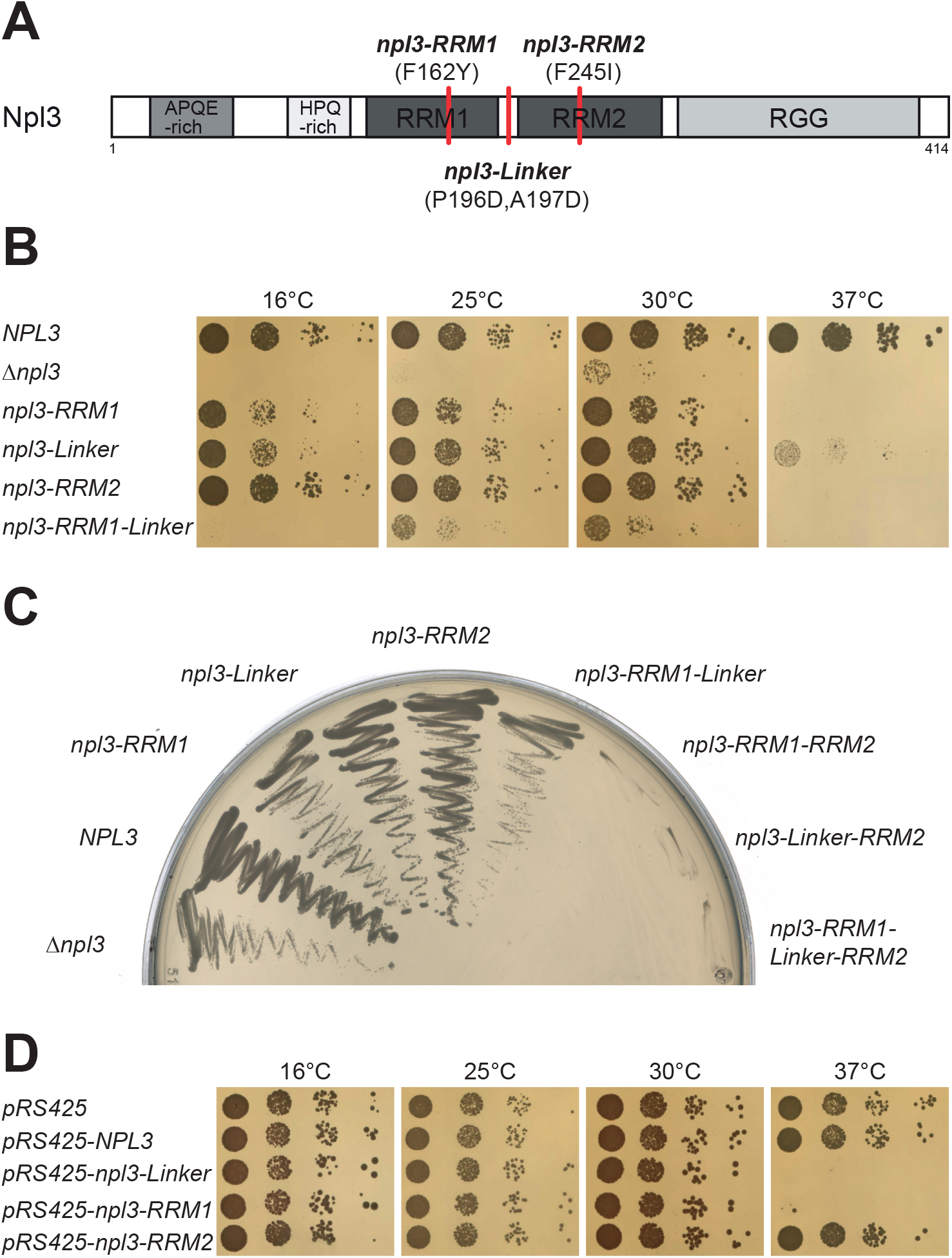
Novel mutations in Npl3 of amino acids binding to RNA. (**A**) Scheme of Npl3 depicting introduced mutations. (**B**) Mutation of amino acids in Npl3 that bind to RNA *in vivo* cause a growth defect. 10-fold serial dilutions of wild-type (wt, *NPL3*), *Δnpl3, npl3-RRM1, -Linker* and *-RRM2* cells were spotted onto YPD plates and incubated for 2-3 days at the indicated temperatures. (**C**) Combination of the *RRM1* and *RRM2* or *Linker* and *RRM2* mutations leads to synthetic lethality. An *NPL3* shuffle strain carrying a genomic deletion of *NPL3* covered by a *URA3*-plasmid encoding *NPL3* was transformed with a plasmid coding for the indicated Npl3 protein and streaked onto an FOA-containing plate to shuffle out the URA3-plasmid and incubated for 3 days at 30°C. And empty plasmid served as comparison for the growth of *Δnpl3* cells. (**D**) Overexpression of npl3-RRM1and npl3-Linker has a dominant negative growth phenotype. 10-fold serial dilutions of wt cells transformed with the high-copy plasmid pRS425 encoding the indicated version of Npl3 were spotted onto SDC(-leu) plates and incubated for 3 days at the indicated temperature.

We first assessed the impact of the mutations on cell growth. Indeed, *npl3-RRM1, -Linker* and *-RRM2* cells display growth defects especially at reduced and elevated temperatures (Figure 2B, Supplementary Figure S2). As expected, the growth defect is less severe than for a complete *NPL3* deletion (Figure 2B, *Δnpl3*). The observed growth defect is not caused by lower Npl3 expression, as the three mutant npl3 proteins are expressed at wild-type (wt) levels (Supplementary Figure S2A).

In addition, we tested whether these three different RNA-binding mutants interact genetically, which is expected if the mutations lead to decreased RNA-binding activity independently of each other. Combination of the RRM1 and the Linker mutations in one protein leads to a strong synthetic growth defect compared to *npl3-RRM1* and *npl3-Linker* cells (Figure 2B, last row). Combination of the RRM1 and RRM2, the Linker and RRM2 as well as all three mutations in one protein leads to lethality (Figure 2C), underlining that the three different RNA-binding mutations have additive effects. This growth defect can be caused by an additive decrease in RNA-binding activity or by the combined defect in binding to different classes of targets by the different domains. In addition, overexpression of the npl3-Linker and npl3-RRM1 protein causes a growth defect at 37°C, a dominant negative effect, which is not observed when wt Npl3 is overexpressed (Figure 1D). Taken together, mutations in amino acids of Npl3 identified to crosslink to RNA *in vivo* lead to a growth defect and display additive effects.

### Both RRM domains and the linker region of Npl3 contribute to RNA binding

We used nuclear magnetic resonance (NMR) spectroscopy to characterize the structure and RNA-binding preference of Npl3 and to assess the effects of the three Npl3 mutations. Three-dimensional structures of the individual RRM domains of Npl3 in the absence of RNA have been previously reported (63,64). However, the mode of RNA recognition by each RRM, the arrangement of the tandem RRM domains (RRM1,2) in the free state or bound to RNA and the RNA-binding surface are poorly characterized. We expressed fragments encompassing the two RRM domains, RRM1-linker-RRM2 (residues 120 to 280, denoted as Npl3^120-280^), of wt Npl3 and the three mutant versions (npl3^120-280^-RRM1 with an F162Y mutation in RRM1, npl3^120-280^-Linker with a P196D/A197D mutation in the linker, and npl3^120-280^-RRM2 with a F245I mutation in RRM2) (Figure 3A) in *E. coli* and analyzed the recombinant proteins by solution NMR (see Supplemental Information).

**Figure 3.**
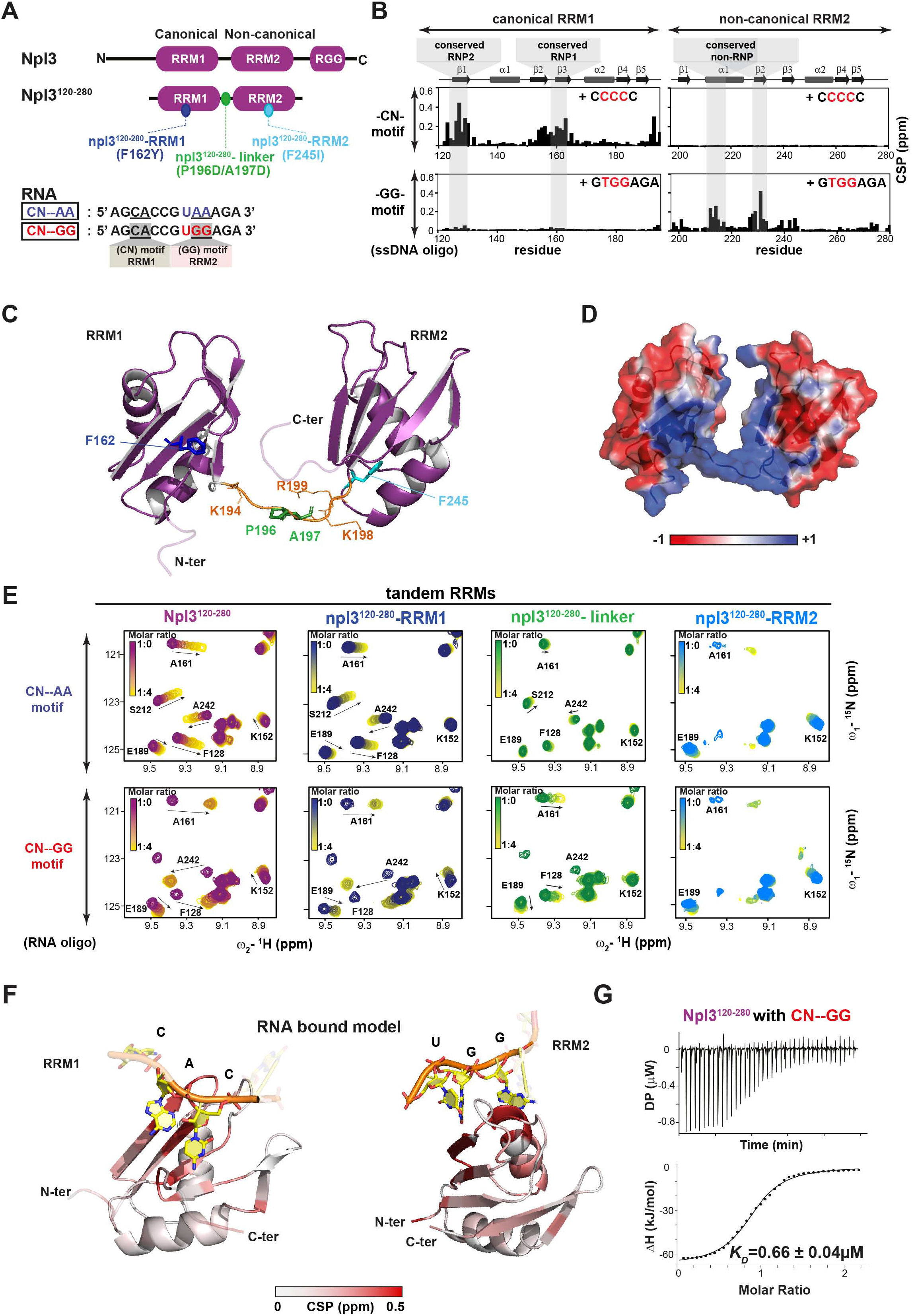
Npl3 RNA binding, structural analysis and effect of mutations. (**A**) Domain organization of full-length Npl3 and the tandem RRM domains used for NMR and binding studies. The position of mutations in npl3-RRM1 (F162Y), npl3-linker (P196D/A197D) and npl3-RRM2 (F245I) are indicated by green, dark-blue and cyan circles, respectively. The two 13-mer RNAs used for binding studies (CN--AA and CN--GG) are indicated below. The key nucleotides in the RNA-binding motifs of RRM1 and RRM2 are underlined. (**B**) NMR chemical shift perturbations (CSPs) with the “CC” and “GG” motifs show the preference of RRM1 (left) and RRM2 (right) for CC and GG, respectively. (**C**) Cartoon representation of the NMR-derived structure of the Npl3 tandem RRM domains. The linker connecting the two RRMs is highlighted in orange. Residues mutated in the linker (P196D/A197D), RRM1 (F162Y) and RRM2 (F245I) are indicated in green, dark blue and light blue respectively. (**D**) Surface representation of the structure colored by electrostatic potential (generated using APBS tool 2.1), blue and red for positive and negative surface charges, respectively. (**E**) Overlay of ^1^H-^15^N NMR correlation spectra of ^15^N-labeled wt RRM1,2 (purple), npl3^120-280^-RRM1 (dark blue), npl3^120-280^-linker (green) and npl3^120-280^-RRM2 (light blue) mutants titrated with “CN--AA” (upper panel) and the “CN--GG” (lower panel) RNA, respectively. Increasing chemical shift changes are colored from purple, green, dark or light blue (free form) to yellow (RNA bound form), respectively. (**F**) NMR and SAXS-derived structural model of the complex of Npl3 with CN--GG RNA where RRM1 and RRM2 recognize a CAC motif (left) and UGG motif (right), respectively. (**G**) ITC binding curves for wt tandem RRM domains of Npl3 with CN--GG RNA ligands are shown.

First, we assessed the RNA-binding preferences for the individual RRM domains of Npl3 using NMR titrations. Single-stranded DNA oligonucleotides were used as first proxies for RNA binding to screen a range of diverse sequences (Figure 3B, Supplementary Figure S4A-F, Supplementary Table S4A). The NMR binding experiments show that cytosine-rich “CC” motif-containing ligands show significant chemical shift perturbation (CSP) at the canonical RNA-binding surface of the RRM1 domain, while no significant binding is observed for the RRM2 domain. In contrast, guanosine-rich “GG” motif-containing ligands strongly bind to RRM2 but not to RRM1 (Figure 3B, Supplementary Figure S4F). In fact, RRM2 has a non-canonical binding interface involving helix α2 and strand β2. This is consistent with the fact that Npl3 shares high sequence conservation with SRSF1 and other SR proteins that exhibit similar features for RRM1 and RRM2 (Supplementary Figure S3A). Taken together, we conclude that RRM1 has a strong preference for “CN”-type motifs, including CCC, while RRM2 prefers “GG” motifs with the UGG motif resembling the RNA-binding preference of the non-canonical RRM in the homologous SRSF1 protein (63,65,66).

Next, we calculated a structural model of Npl3’s tandem RRM domains based on NMR and small angle X-ray scattering (SAXS) data (Figure 3C, Supplementary Figure S5A-E, Supplementary Tables S5 and S6). The structural model shows that the β-sheet surfaces of the two RRM domains face towards each other, forming a positively charged surface (Figure 3D). The two domains adopt a rather fixed orientation, and the linker connecting the two RRM domains has only reduced flexibility (Supplementary Figure S3B-E). Interestingly, the three *npl3* mutations tested *in vivo* map to the positively charged region including the linker. We next evaluated the structural integrity of RRM1-Linker-RRM2 fragment containing the different mutations by comparing their NMR spectra with the wt version (Supplementary Figure S6A). For the RRM1 mutant (npl3^120-280^-RRM1), no significant spectral changes are observed compared to wt, suggesting that F162Y does not affect the fold of the tandem RRM domains (Supplementary Figure S6 A, B, right panel). For the npl3^120-280^-Linker mutant, notable spectral changes are observed only in the proximity of the linker (Supplementary Figure S6B, middle panel), indicating that the overall structure is largely unperturbed. However, comparison of the SAXS data for Npl3^120-280^ and npl3^120-280^-Linker (Supplementary Figure S6C, Supplementary Table S5) indicates that the RRM1,2 domain arrangement is slightly more compact in the linker mutant, perhaps as a consequence of replacing the more rigid and extended P196 residue. In contrast, the F245I mutation in RRM2 (npl3^120-280^-RRM2) causes severe line-broadening for most residues in RRM2, while the NMR signals for the RRM1 are largely unaffected (Supplementary Figure S6 A, B, right panel). Overall, we conclude that npl3^120-280^-RRM1 and npl3^120-280^-Linker maintain the overall structure of the tandem RRM domains, while npl3^120-280^-RRM2 strongly affects the RRM2 fold and is thus expected to impair protein functions involving RRM2.

To characterize the RNA-binding activity of the tandem RRM domain and the role of the linker, we analyzed the interaction with two RNA sequences, which harbor the respective binding motifs of RRM1 (CN) and RRM2 (GG). Specifically, we tested binding to CN--GG (5’-AG**<u>CA</u>**CCG**U<u>GG</u>**AGA-3’) and a variant CN--AA (5’-AG**<u>CA</u>**CCG**U<u>AA</u>**AGA-3’), where RNA binding by RRM2 is expected to be strongly reduced. To this end, we performed NMR and isothermal titration calorimetry (ITC) experiments using the two RNA oligonucleotides with a ^15^N-isotopically labelled RRM1-Linker-RRM2 fragment (Npl3^120-280^). The CN--AA oligo binds to both RRM domains with modest affinity as NMR signals shift with increasing concentration of the RNA ligand, and saturation is obtained only at 4-fold molar excess (Figure 3E, Supplementary Figure S7). CSPs are observed for both RRM domains and the linker region, indicating that this RNA interacts with both RRM domains, and the canonical RNP motifs in RRM1 are most strongly affected (Supplementary Figure S8A). The positively charged residues K194, K198 and R199 in the linker also show significant perturbation, which demonstrates that the linker contributes to RNA binding. The interaction of Npl3^120-280^ with the CN--AA RNA was not detectable in ITC experiments indicating weak binding (Supplementary Figure S8C, left panel, Supplementary Table S4B). This is confirmed by NMR titration experiments, which show an average *K*_*D*_ for the interaction of *K*_*D*_ ≈ 150 μM (Supplementary Figure S8C, right panel). In contrast, the binding to the CN--GG RNA is significantly stronger, consistent with NMR titrations showing binding kinetics in the intermediate to slow exchange regime (Figure 3E, Supplementary Figure S7A, right panel). Spectral changes map to the same binding surface seen for the CN--AA RNA, but are much stronger and involve more residues of RRM2 (Supplementary Figures S7A, S8A). ITC experiments show high affinity binding of the CN--GG RNA with Npl3^120-280^ with *K*_D_ = 0.66 µM (Figure 3G, Supplementary Figure S8C, Supplementary Table S4B). Interestingly, the RNA-binding region maps to the β-sheets of the canonical RNP sites in RRM1, the non-canonical conserved regions in RRM2 and the positively charged surface in Npl3^120-280^ (Figure 3D, Supplementary Figure S3A).

We then calculated a structural model of the Npl3^120-280^ complex with the high-affinity CN--GG RNA ligand based on NMR and SAXS data (Figure 3F, Supplementary Figure S5F,J,K), which rationalizes the RNA-binding features observed. The overall RRM domain arrangement is very similar to the one observed in the absence of RNA (Supplementary Figure S5G). As expected from the sequence conservation, the RNA-binding interfaces and interactions of the RRM domains are very similar to the corresponding interactions of SRSF1 (Supplementary Figure S5I).

Next, we investigated the effects of the Npl3 mutations on RNA binding (Figure 3E). A strongly reduced RNA binding of the linker mutation may be rationalized by charge clashes of two aspartate residues in the mutant linker. This may also contribute to the small but significant change in the RRM domain distances (as seen by SAXS, Supplementary Figure S6C, Supplementary Table S5), which may reduce optimal contacts with the RNA. To further characterize the effects of the linker mutation, we titrated both RNA ligands to the npl3^120-280^-Linker mutant fragment. Virtually no spectral changes are seen at 4-fold excess of the CN--AA RNA, while a strongly reduced interaction is observed with the CN--GG RNA (Figure 3E, Supplementary Figures S7C, S8A). Consistent with this, no binding is detected in ITC experiments (Supplementary Figure S8B,C, middle panel, Supplementary Table S4B). Taken together, our data demonstrate that the linker mutation does not affect the structural integrity, but strongly reduces RNA-binding affinity. The npl3^120-280^-RRM1 mutant shows reduced RNA binding with *K*_D_ = 10 μM for the CN--GG RNA, thus 15-fold reduced compared to wt Npl3^120-280^ (Figure 3E, Supplementary Figures S7B, S8A, B, Supplementary Table S4B). The npl3^120-280^-RRM2 protein (in which RRM2 is destabilized, Figure 3E, Supplementary Figure S6A right panel, Supplementary Figure S7D) shows a 10-fold reduced RNA-binding affinity (*K*_D_ = 5 μM) for the CN--GG RNA in ITC experiments, compared to wt Npl3^120-280^ (Supplementary Figure S8B, right panel, Supplementary Table S4B). In summary, the point mutations in the RRM1, linker or RRM2 regions of Npl3 strongly reduce the RNA-binding activity compared to wt Npl3^120-280^ for both RNA sequences tested.

To assess whether RNA binding of the npl3 mutants is also decreased in the context of the full-length protein *in vivo*, we performed RNA immunoprecipitation (RIP) experiments. Genomically TAP-tagged Npl3 variants were immunoprecipitated from whole-cell extracts. Co-immunoprecipitated RNA was analyzed by reverse transcription (RT) and quantitative PCR (qPCR) for three representative transcripts (*PMA1, CCW12* and *YEF3*). All three mutant Npl3 proteins bind to these mRNAs significantly less than the wt protein (Figure 4A). Thus, mutations of the amino acids crosslinked to RNA that lead to a growth defect indeed reduce the RNA-binding activity of Npl3. In line with our *in vitro* assays, the effects tend to be strongest for the *npl3-Linker* mutation, underlining that the short linker region seems to play an important role for the RNA-binding activity of Npl3.

**Figure 4.**
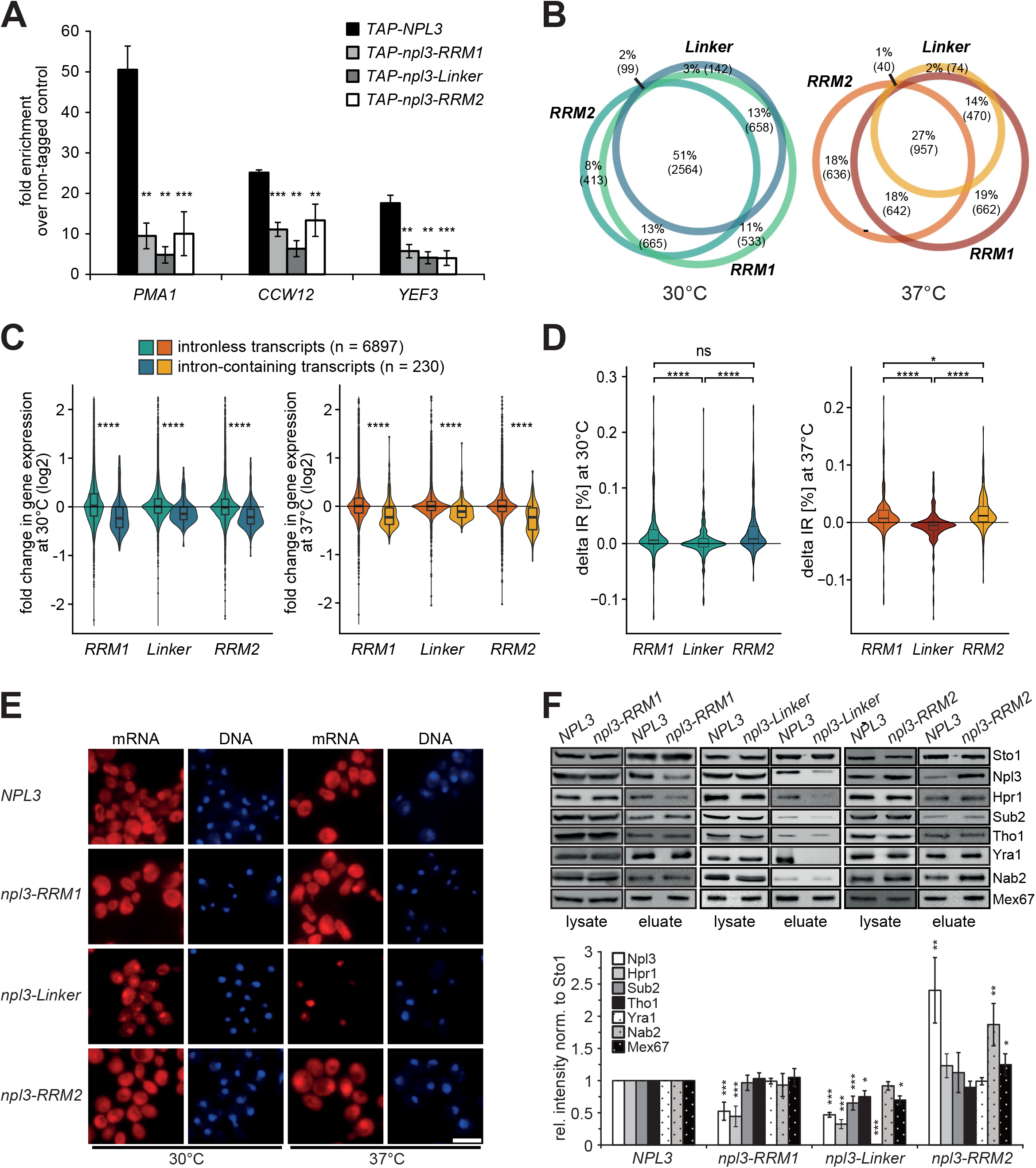
Mutations in *NPL3* that decrease RNA-binding activity cause different effects on nuclear mRNA export, nuclear mRNP composition and gene expression. (**A**) The three Npl3 mutants bind less mRNA *in vivo*. TAP-tagged wt and mutant versions of Npl3 were immunoprecipitated from whole-cell extract and bound RNA quantified by RT-qPCR for the indicated transcripts. The amount of co-immunoprecipitated mRNA was calculated as the enrichment over the amount of RNA in a control experiment with a strain expressing non-tagged Npl3. (**B**) *npl3* mutants show similar changes in transcript abundance. Venn diagrams depicts overlap of differentially expressed transcripts (adjusted p-value < 0.05) in the three mutants at 30°C (left) and after shift to 37°C for one hour (right). (**C**) Intron-containing transcripts show a significant trend for downregulation. Violin boxplots compare log_2_-transformed fold changes in transcript abundance for intron-containing (n = 230) and intron-less (n = 6,897) transcripts in the three *npl3* mutants compared to wt at 30°C (left) and 37°C (right). **** p-value ≤ 0.0001, Wilcoxon rank-sum test. (**D**) Introns show increased retention in the *npl3-RRM1* and *npl3-RRM2* mutants. Beeswarm-boxplots compare difference in intron retention (ΔIR; n = 215) for each *npl3* mutant in relation to wt at 30°C and 37°. * p-value < 0.05, **** p-value ≤ 0.0001, Wilcoxon rank-sum test. (**E**) *npl3* mutant cells show different degrees of an mRNA export defect. The localization of bulk mRNA was visualized by *in situ* hybridization with fluorescently labeled oligo(dT) in *npl3* mutant cells grown at 30°C or shifted to 37°C for one hour. DNA was stained with DAPI. (**F**) Nuclear mRNP composition changes in the three different *npl3* mutants. Western blots (upper panel) and quantification of three independent experiments (lower panel) of nuclear mRNPs purified via Cbc2-TAP purification from wt, *npl3-RRM1, npl3-Linker* and *npl3-RRM2* cells. The amounts of Sto1, Npl3, Hpr1, Sub2, Tho1, Yra1, Nab2 and Mex67 were quantified and normalized to the amount of the CBC subunit Sto1. Values for wt cells were set to 1.

### *npl3* mutations globally affect transcript abundance and splicing

Based on our initial findings, we decided to globally investigate the transcriptome of the three *npl3* mutants. To this end, we harvested RNA from wt and mutant cells and performed polyA^+^ RNA sequencing (RNA-seq). Cells were either grown at 30°C or shifted to 37°C for one hour because of the small but marked growth difference between wt cells and all three mutant cells at this time point (Supplementary Figure S2C, E, G). Differential expression analysis using DESeq2 (56) reveals widespread changes (adjusted p-value < 0.05; Supplementary Figure S9A, Supplementary Table S7). At 30°C, for each *npl3* mutant more than 3,500 transcripts differ significantly in abundance, which are mostly shared between mutants (Figure 4B, left panel). In contrast, at 37°C, less transcripts differ in total, but the three mutants separate more clearly (Figure 4B, right panel). At both temperatures, the *npl3-RRM1* and *npl3-Linker* mutants show a similar trend, whereas the *npl3-RRM2* mutant is set apart (Supplementary Figure S9B). Taken together, changing the RNA-binding activity of Npl3 leads to significant transcriptome alterations.

We next investigated differences in splicing. As a first hint, we observed that the abundance of intron-containing transcripts is significantly reduced in all three *npl3* mutants, most prominently in the *npl3-RRM1* and *npl3-RRM2* mutants at 37°C (Figure 4C), suggesting that splicing is affected. We therefore used the RNA-seq reads at intron-exon boundaries to quantify the splicing efficiency. Indeed, intron retention is elevated in the *npl3-RRM1* and *npl3-RRM2* mutants at 37°C (Figure 4D). In contrast, the *npl3-Linker* mutant displays an opposing pattern, such that intron retention is reduced at 37°C. Thus, the mutations in *NPL3* affect splicing efficiency in opposing ways. Taken together, mutations in *NPL3* result in profound alterations of the transcriptome. The three mutants show overlapping, but distinct phenotypes, likely reflecting divergent functions of the RRM and linker domains of Npl3, which might be mediated by different RNA-binding preferences of the individual RNA-binding domains.

### *npl3* mutations cause distinct effects on nuclear mRNA export and nuclear mRNP composition

To assess the functional consequences of the three mutations, we next tested nuclear mRNA export efficiency. We performed RNA fluorescence *in situ* hybridization (RNA-FISH) against poly(A) tails to visualize the global mRNA distribution. Interestingly, even though the RNA-binding activity is reduced in each of the three mutants, they show different phenotypes (Figure 4E). Whereas *npl3-RRM1* cells have no mRNA export defect at 30°C and a minor defect at 37°C, *npl3-Linker* cells already display a strong mRNA export defect at 30°C, which is exacerbated at 37°C. *npl3-RRM2* cells show an intermediate phenotype with a mild defect at 30°C and a stronger defect at 37°C (Figure 4E). Interestingly, the strength of the mRNA export defect does not correlate with the severity of the growth defect (Figure 2B), indicating that other processes than mRNA export are probably impaired in these mutants.

As nuclear mRNA export is decreased in the *npl3* mutants, we determined whether the composition of nuclear mRNPs is changed in these cells. Nuclear mRNPs were purified by native purification of endogenously TAP-tagged Cbc2, and the amount of co-purifying Npl3 as well as six other nuclear mRNP components was determined by Western blot. These encompass the THO/TREX subunits Hpr1, Sub2 and Yra1, the nuclear poly(A)-binding protein Nab2, Tho1 and the mRNA exporter subunit Mex67 (Figure 4F, upper panel). Importantly, the *in vivo* RNA-binding activity of Cbc2 in *npl3-RRM1* and *npl3-Linker* cells is similar to wt as equal amounts of three representative mRNAs copurify with Cbc2 in a RIP experiment (Supplementary Figure S10). Consequently, also comparable amounts of nuclear mRNPs co-purify with Cbc2 in these two mutants, in contrast to the *npl3-RRM2* mutant. As expected from its decreased RNA-binding affinity, the amount of Npl3 co-purifying with nuclear mRNPs is decreased in the *npl3-RRM1* and *npl3-Linker* mutants (Figure 4F). Regarding the other six mRNP components, only the amount of Hpr1 is decreased in the *npl3-RRM1* mutant, which is consistent with the very mild mRNA export defect (Figure 4F). In contrast, the abundance of Hpr1, Sub2, Tho1 and Mex67 is decreased in nuclear mRNPs of *npl3-Linker* cells (Figure 4E). Thus, Npl3 is needed for the recruitment to or retention of Hpr1, Sub2, Tho1 and Mex67 at the nuclear mRNP. This change in the composition of nuclear mRNPs is consistent with the mRNA export defect observed in *npl3-Linker* cells (Figure 4E). Despite the decreased RNA-binding activities of Npl3 and Cbc2 in these cells, the relative amount of Npl3 in nuclear mRNPs is increased in the *npl3-RRM2* mutant (Figure 4E), potentially due to unfolding and aggregation of RRM2 induced by the mutation. In addition, the amount of Nab2 and Mex67 in nuclear mRNPs is also specifically increased in *npl3-RRM2* cells (Figure 4E). Taken together, the three *npl3* mutants impact nuclear mRNA export and nuclear mRNP composition differently, with the strongest effects observed for the *npl3-Linker* mutant. This underlines the notion that – despite the fact that mRNA binding is decreased in each of these mutants – they have different effects on gene expression processes, possibly by divergent RNA sequence preferences of the three RNA-binding sites.

### *npl3-Linker* cells show changes in nuclear mRNP composition

Based on its strong nuclear mRNA export defect, the differences in nuclear mRNP composition and the divergent splicing changes, we chose the *npl3-Linker* mutant for further analysis. To assess the changes in nuclear mRNP composition in an unbiased way, we purified Cbc2-TAP from wt and *npl3-Linker* cells and used a label-free MS quantification approach to assess abundances of the co-purified proteins (Supplementary Table S8). Consistent with the results obtained by Western blot experiments, the *npl3-linker* mutation results in the prominent reduction of the intensities of Npl3 itself as well TREX complex components (Figure 5A). The intensities of Tho1 and the mRNA exporter subunits Mex67 and Mtr2 are also decreased, although they do not satisfy the significance criteria of FDR<2% (Figure 5A). In addition, reflecting the involvement of Npl3 in splicing, ribosome biogenesis and translation, the intensities of many splicing factors, ribosome biogenesis factors, ribosomal proteins and translation factors are also significantly decreased (Figure 5A). In summary, with the exception of Nab2, the amount of nuclear mRNP components present in nuclear mRNPs decreases in *npl3-Linker* cells.

**Figure 5.**
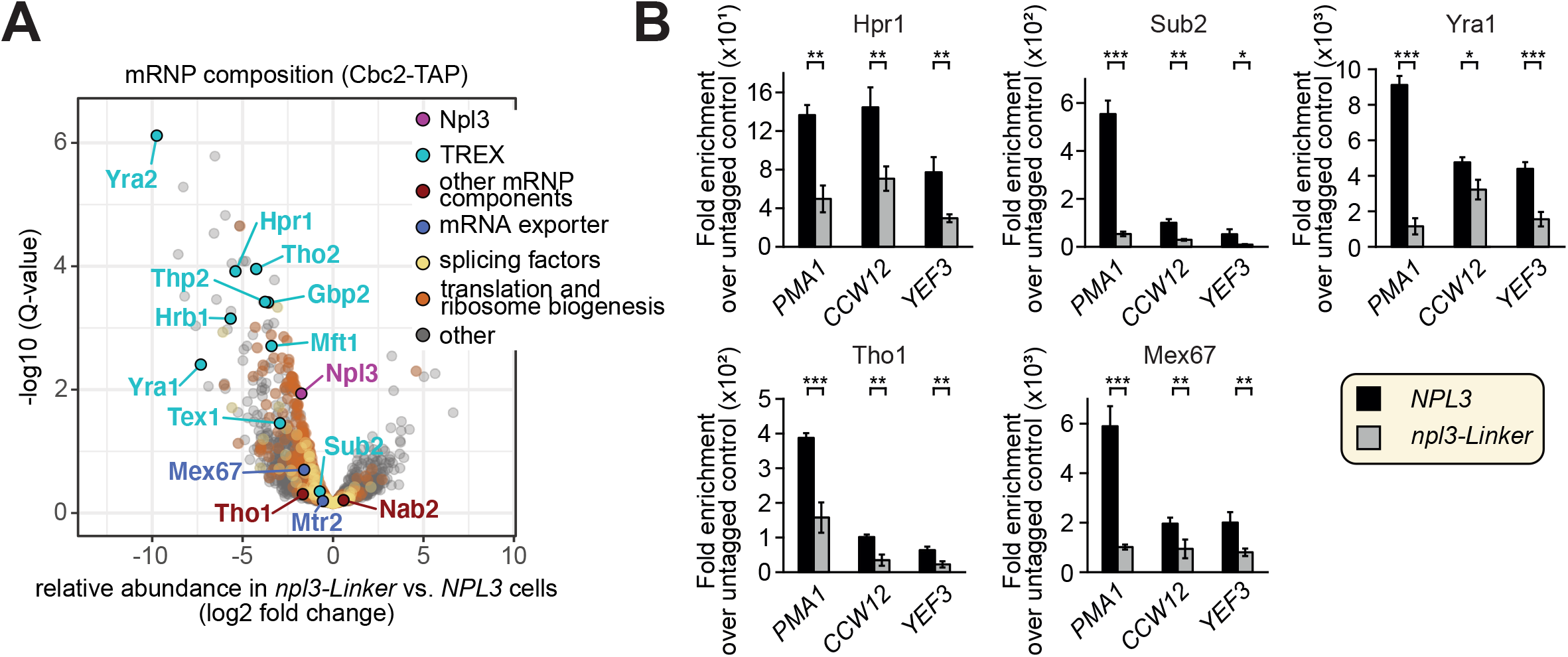
*npl3-Linker* cells show changes in nuclear mRNP composition, gene expression and splicing. (**A**) Mass spectrometric analysis of changes in the composition of nuclear mRNPs in *npl3-Linker* cells. The Volcano plots depicts the fold change (x-axis) and statistical significance (y-axis) of all proteins detected by MS. Npl3 is shown in purple, TREX components in blue, the mRNA exporter in light blue and further mRNP components in green, and all identified proteins are labeled with their corresponding names. Splicing factors are depicted in brown and ribosome biogenesis factors, ribosomal proteins and translation factors in dark grey. All other proteins are shown in light grey. (**B**) mRNA binding of Hpr1, Sub2, Tho1 and Mex67 is decreased in *npl3-Linker* cells *in vivo*. RIP experiments of Hpr1, Sub2, Tho1 and Mex67 as in Figure 4A.

The decrease in the amount of TREX complex, Tho1 and Mex67 in nuclear mRNPs suggests that these proteins bind less mRNA *in vivo* in the *npl3-Linker* mutant. To assess this, we performed RIP experiments with these proteins from *npl3-Linker* cells. Indeed, the TREX components Hpr1 and Sub2, Tho1 and the mRNA exporter subunit Mex67 bind less mRNA as assessed for three representative transcripts (Figure 5B). Thus, Npl3 function is required for stable association of these nuclear mRNP components with nuclear mRNPs.

### Npl3, Hpr1, Yra1 and Mex67 occupancy at transcribed genes is decreased in *npl3-Linker* cells

Nuclear mRNP components are recruited to the mRNA already during transcription. Thus, the decrease in some of the nuclear mRNP components could be due to a decrease in their recruitment to transcribed genes. As this recruitment depends on the presence of RNA and thus on transcription, we first assessed the occupancy of RNA polymerase II (RNAPII) at three representative genes, *PMA1, CCW12* and *YEF3*, by chromatin immunoprecipitation (ChIP) (Figure 6A). RNAPII occupancy is increased, whereas the occupancy of Npl3 is decreased in *npl3-Linker* compared to wt cells (Figure 6B, C, D). Based on its RNA-dependent recruitment to transcribed genes, this decrease is probably due to the reduced RNA-binding activity of the npl3-Linker protein. Consistent with its decrease in nuclear mRNPs, the occupancy of the THO component Hpr1 decreases in the *npl3-Linker* mutant (Figure 6E), suggesting that recruitment to or retention of THO at transcribed genes depends on Npl3. The occupancy of the TREX component Yra1 and the mRNA exporter subunit Mex67 at transcribed genes also decreases in *npl3-Linker* cells (Figure 6H, I). In contrast, the occupancies of Sub2 and Tho1 on chromatin do not change (Figure 6F, H). Thus, the decreased presence of the nuclear mRNP components Hpr1, Yra1 and Mex67 versus Sub2 and Tho1 in mRNPs appears to have different causes: Whereas Hpr1, Yra1 and Mex67 fail to be efficiently recruited to or retained at transcribed genes, Sub2 and Tho1 are not efficiently transferred from the site of transcription onto the forming mRNP in *npl3-Linker* cells. The RNA-binding activity of Npl3 is required for both processes. In summary, we identify two novel functions of the SR-like protein Npl3 based on the identification and mutation of RNA-binding sites.

**Figure 6.**
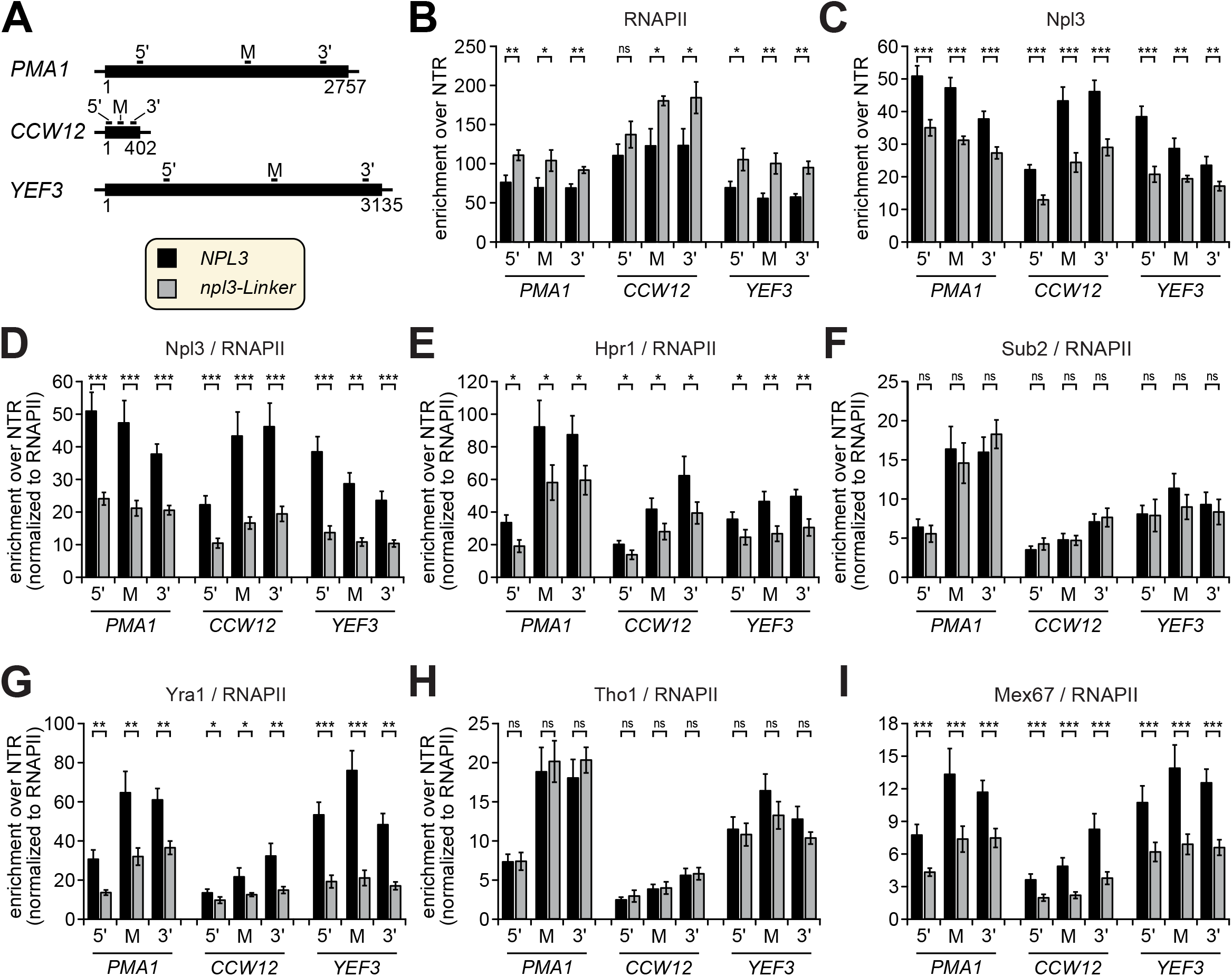
The occupancy of Npl3, Hpr1 and Mex67 at transcribed genes is decreased in *npl3-Linker* cells. (**A**) Scheme of the three exemplary genes *PMA1, CCW12* and *YEF1*. Open reading frames (ORFs) are represented by a solid line. The bars above the genes indicate the position of the primer pairs used for analysis of the ChIP experiments. Numbers indicate the nucleotides of each gene. 5’: 5’ end of the ORF, M: middle, 3’: 3’ end. (**B**) The occupancy of RNAPII is increased in *npl3-Linker* cells. (**C and D**) The occupancy of Npl3 (B) and the occupancy of Npl3 normalized to the occupancy of RNAPII (D) is decreased in *npl3-Linker* cells. (**E**) The occupancy of Hpr1 normalized to RNAPII is decreased. (**F and G**) The occupancy of Sub2 (F) and of Tho1 (G) normalized to RNAPII is unchanged. (**H**) The occupancy of Mex67 normalized to RNAPII in *npl3-Linker* cells is decreased.

## DISCUSSION

Every RNA metabolic process entails a multitude of RNA-protein interactions. Affinity purification of proteins crosslinked to RNA is frequently used to identify the crosslinked RNA and the site(s) of its interaction with the protein (67-71). Here, we identified for the first time RNA interaction sites on proteins at amino acid resolution after affinity purification of tagged proteins with their crosslinked RNA isolated from UV-crosslinked cells. RNA-protein UV-crosslinking combined with MS is the method of choice for the identification of RNA-binding proteins and their binding regions. In earlier *in vivo* crosslinking approaches, crosslinked RNA moieties were isolated (e.g. through their poly(A) tail or an aptamer) and the crosslinked proteins were then examined by MS to obtain information about their RNA-binding regions and the crosslinked amino acids (5,8,59-61). We successfully applied the converse method, *i*.*e*. specific affinity purification of the crosslinked protein moiety, and we attribute its success to the sensitive and very specific enrichment of crosslinked peptide-oligoribonucleotide conjugates (5) in combination with the sensitive MS instrumentation and the software tool RNP^XL^ and its successor NuXL for improved MS data evaluation and annotation ((7) and unpublished, respectively).

Our data allow us to map the crosslinking sites on available 3D structures of the THO-Sub2 complex from yeast containing the proteins Tho2, Hpr1, Thp2, Mft1, Tex1 and Sub2, but no RNA (72-74), and this in turn gives a picture of how RNA interacts with the proteins Tho2, Hpr1, Mft1 and Sub2 (Supplementary Figure S11).

Our approach can be adapted to any cellular system, and it might be interesting to combine it directly with CLIP methods and its derivatives (67). This may be expected to yield complementary information on the crosslinked RNA and the crosslinking sites within the protein of interest. The fact that we are able to pinpoint the crosslinked peptide/amino acid experimentally makes our approach feasible for high-throughput studies that aim to obtain an inventory of RNA-binding peptides or amino acids of particular RBP complexes. This will be a valuable source for functional studies by e.g. mutagenesis of experimentally determined RNA-binding peptides / amino acids. In fact, the MS-based identification of RNA-crosslinking sites in RRM1 (F162), the linker region (P196, A197) and RRM2 (F229, S230) of Npl3 enabled us to generate mutant proteins and to study the effect of these mutations on the structure of Npl3 and on its function in mRNP biogenesis.

The RRM and RGG domains of Npl3 are known to have RNA-binding activity (63,75). Indeed, RRM1 adopts a canonical fold. In contrast, RRM2 lacks the conserved RNP1 and RNP2 motifs and instead features a conserved sequence motif “SWQDLKD” in helix α1, characteristic of a non-canonical, so-called pseudo-RRM domain (63,64,66). Here, we present a structural model for the tandem RRM domains that shows a large positively charged surface of Npl3 including the linker connecting the two RRM domains.

Our RNA-binding studies show distinct binding preferences for the two RRM domains. The canonical RRM1 recognizes a CC motif and the non-canonical RRM2 shows specificity for a GG motif. Accordingly, the binding affinity of the tandem RRM1,2 domains to CN--AA RNA (N being any nucleotide) is significantly lower (150-fold) compared to a CN—GG RNA ligand. Interestingly, the AG**<u>CAC</u>C**G**<u>UGG</u>**AGA RNA binds to RRM1-RRM2 with 5’ to 3’ orientation, different to most other tandem RRM domains. The two RRM domains are partially prearranged for RNA binding, and, consistently, the overall domain arrangement of the free and RNA-bound tandem RRM domains is relatively similar. The short linker is flexible in the absence of RNA, potentially allowing fine-tuning of the domain arrangement to optimize RNA recognition. Npl3 is homologous to human SRSF1, and our analysis indeed confirms that the RNA-binding preferences for RRM1 and RRM2 are comparable to human SRSF1 (65,66).

Surprisingly, our data show that the linker connecting RRM1 and RRM2 strongly contributes to RNA binding (Supplementary Figure S8A). Notably, the overall reduction in RNA-binding affinity upon linker mutation observed *in vitro* correlates well with effects observed *in vivo*. Likely, the increased flexibility by replacing the proline and charge reversal by two aspartates in this region contribute to a reduction in RNA-binding affinity. Interestingly, the RRM1 mutation also reduces RNA binding by 15-fold compared to the wt protein, suggesting an important role for the tyrosine hydroxyl and altered stacking interactions. The mutation in RRM2 leads to domain unfolding, which rationalizes the significantly reduced binding affinity (∼7 fold) to CN--GG RNA. Interestingly, no well-defined RNA-binding motif has been identified *in vivo* (76). This may reflect that differential contributions of the two tandem domains of Npl3 may enable binding to distinct substrates depending on the process. Taken together, our NMR and biochemical data demonstrate a critical role of the Npl3 linker, the conserved sequence motifs in RRM1 and the non-canonical RRM2 for RNA binding.

Although mRNA binding of all three mutant proteins is reduced *in vitro* and *in vivo* (Figure 3, 4A), the functional consequences differ. Growth of the three *npl3* RNA-binding mutants is affected to a varying extent, likely resulting from the sum of the different processes impaired in these mutants. For example, splicing efficiency, nuclear mRNP composition and mRNA export are differentially affected in the three mutants. Npl3 is the only SR-like protein that promotes splicing in *S. cerevisiae* through co-transcriptional recruitment of the U1 and U2 snRNPs to chromatin (77). Consistently, all three *npl3* mutants are defective in splicing. However, while deletion of *NPL3* results in intron retention in a subset of pre-mRNAs (77), which also occurs in the *npl3-RRM1* and *npl3-RRM2* mutants (Figure 4D), intron retention decreases in the *npl3-Linker* mutant, suggesting that mRNAs are more efficiently spliced. This may reflect the longer residence time of mRNA in the nucleus of *npl3-Linker* cells as a consequence of the strong nuclear mRNA export defect in this mutant compared to the other two.

We selected the *npl3-Linker* mutant for a detailed analysis based on this strong nuclear mRNA export defect (Figure 4E). It is possible that – in addition to the reduced RNA binding – the *npl3-Linker* mutation also abrogates or decreases protein-protein interactions. Thus, the question remains whether the decrease in RNA-binding activity is the cause for the defects observed in the *npl3-Linker* mutant. The interaction of Npl3 with most, if not all, mRNP components depends at least partially on RNA, as RNase treatment decreases the co-purification of Sto1, Sub2, Yra1, Mex67, Tho1 and Nab2 (data not shown). The RNA-independent protein interactions, however, are largely unchanged between npl3-Linker and Npl3 (data not shown), suggesting that protein-protein interactions are not significantly impaired in the *npl3-Linker* mutant and supporting the interpretation that loss of RNA binding plays a causal role in the mutant phenotypes. In this mutant, RNAPII occupancy across protein-coding genes is increased (Figure 6B), either as a compensatory response of the cells to increase mRNA production in the face of the compromised downstream processes, or reflecting increased RNAPII stalling caused by inadequate mRNA processing, packaging and nuclear export. According to the current model, mRNP components are recruited to the nascent mRNA in a co-transcriptional manner and transferred to the mRNA when it leaves the site of transcription. Interestingly, the occupancy of Hpr1 and thus most likely the whole THO complex, Yra1 and Mex67 at protein-coding genes is decreased in *npl3-Linker* cells (Figure 6). Consistently, these proteins also have a lower abundance in nuclear mRNPs (Figures 4F, 5A, B). In contrast, the occupancy of Sub2 and Tho1 is not affected in *npl3-Linker* cells (Figure 6F and H), but their level in nuclear mRNPs is decreased (Figure 4F, 5A, B). Thus, Npl3 could either be required for the transfer of Sub2 and Tho1 from the transcription machinery onto the mRNA or for their retention within the mRNP. These decreases are specific for particular mRNP components as the amount of Nab2 in nuclear mRNPs is not affected in the *npl3-Linker* mutant (Figure 4F). Importantly, this specific RNA-binding mutant of Npl3 allowed us to reveal novel functions of Npl3 in promoting the occupancy of THO, Yra1 and Mex67 at transcribed genes and for the presence of these three mRNP components as well as Sub2 and Tho1 in nuclear mRNPs.

This is the first time that Npl3 function has been linked to Sub2 and Tho1. The human orthologs of Sub2, Tho1 and Yra1 are UAP56/DDX39B, SARNP and ALYREF, respectively, which form an ATP-dependent trimeric complex (27). We show here that Npl3 function is needed for the transfer of Sub2 and Tho1 from the site of transcription onto nuclear mRNPs and/or their retention therein (Figure 4F). Interestingly, Sub2 and Tho1 require each other for their association with nuclear mRNPs, consistent with their direct binding to each other (data not shown). In contrast, they are not needed for the presence of Npl3 in nuclear mRNPs (data not shown). Thus, Npl3 is required for Sub2 and Tho1 in nuclear mRNPs but not vice versa. Furthermore, the decrease of Mex67 levels in nuclear mRNPs in *npl3-Linker* cells (Figure 4F) is consistent with the observed nuclear mRNA export defect (Figure 4E), which is most likely also caused by the decrease in other mRNP components and thus might be part of a quality control mechanism.

In summary, the identification of RNA-binding sites within RBPs, here the nuclear mRNP components, and the detailed analysis of specific RNA-binding mutants of Npl3 uncovered two novel functions of this protein in mRNP biogenesis. Npl3 is necessary to recruit the mRNP components Hpr1, Yra1 and Mex67 to the site of transcription and to transfer and/or retain the mRNP components Sub2 and Tho1 at nuclear mRNPs. This underscores the utility of a targeted approach for the identification and mutational analysis of amino acids binding to RNA to unravel the function of RNA-binding proteins implicated in complex RNA processing networks.

## AVAILABILITY and ACCESSION NUMBERS

The MS protein-RNA crosslinking data (Figure 1, Supplementary Tables S1, S2, S3) were deposited to the ProteomXchange repository with the dataset identifier PXD035153. The MS proteomic data (Figure 5A, Supplementary Table S8) were deposited to the ProteomeXchange repository with the dataset identifier PXD034656.

The RNA-seq data generated in this study is available in the Gene Expression Omnibus (GEO) under the accession number GSE160709.

## SUPPLEMENTARY DATA

Supplementary Data are available at NAR online.

## ACKNOWLEDGEMENT

We thank Vladislav Kunetsky for graphical design of the model shown in Figure 7 and Cornelia Kilchert (Institute of Biochemistry, FB08, JLU) for critical reading of the manuscript. We are thankful to Susanne Röther for the *NPL3* shuffle strain and plasmid pRS315-*NPL3* and to Britta Coordes for the *TAP-NPL3* strain. We thank Sabine König, Ralf Pflanz, Uwe Plessmann and Monika Raabe (MPI-NAT, Göttingen) for assistance in MS. We thank Sam Asami and Gerd Gemmecker for support with NMR experiments. We acknowledge SBGrid and the NMRbox server (78) for providing access to NMR and structure calculation softwares. We acknowledge NMR measurements at the Bavarian NMR Center (BNMRZ) and SAXS measurements at the facility of the SFB1035 at Department Chemie, Technical University of Munich, Germany. The antibody directed against Cbp80 was a kind gift of Dirk Görlich (MPI Goettingen, Germany) (36).

**Figure 7.**
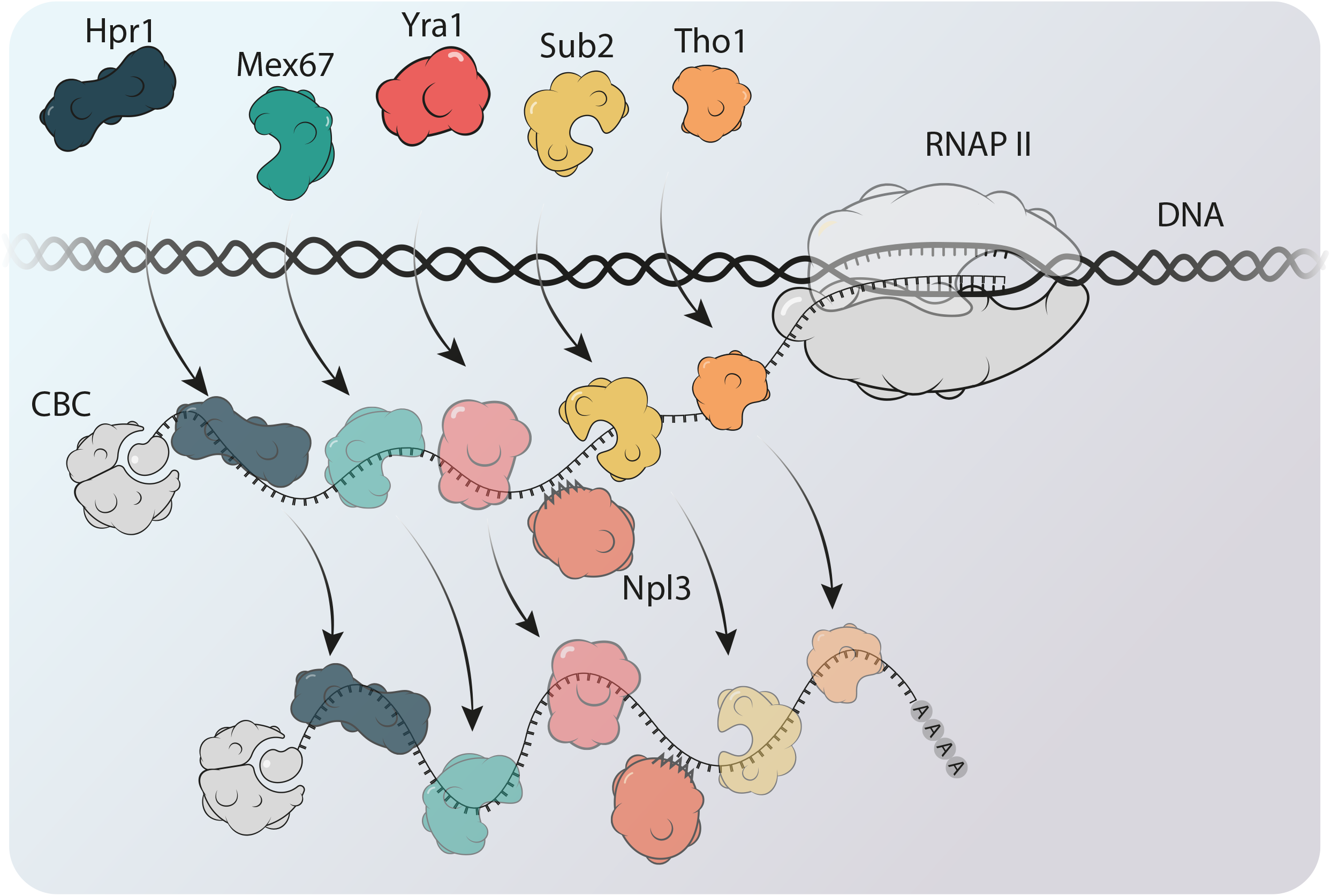
Model of Npl3 function in nuclear mRNP assembly. Npl3 has two functions in nuclear mRNP assembly. Npl3 is needed for the recruitment and/or retention of Hpr1 – and thus most likely the THO complex – as well as Yra1 and Mex67 to transcribed protein-coding genes. In addition, Npl3 transfers the mRNP components Sub2 and Tho1 from the site of transcription to the mRNA. Impairment of the RNA-binding activity of Npl3 causes a decrease of these mRNP components in nuclear mRNPs.

