## Supplementary Data for "Npl3 functions in mRNP assembly by recruitment of mRNP components to the transcription site and their transfer onto the mRNA"

### Design of Npl3 mutants with impaired RNA-binding activity

In order to obtain Npl3 mutants with reduced RNA-binding activity, we mutated amino acids in the RNA recognition motif 1 (RRM1), linker and RRM2 domains of Npl3 that cross-linked to RNA *in vivo* (Supplementary Table S1 and Supplementary Figure S1A). For RRM1 we tested single, double and triple mutations of the three conserved phenylalanines (F) to tyrosine (Y) in the RNP1 motif (F160, F162 and F165; Supplementary Figure S1A). All mutant npl3 proteins are expressed at wild-type (wt) levels *in vivo* (Supplementary Figure S1B). Since mutation of F162 is sufficient for the observed growth defect and the temperature sensitive phenotype (ts and cs, Supplementary Figure S1C), we chose this mutant for further analysis. In the linker region we mutated proline (P) 196 and alanine (A) 197 to aspartic acid (D). In RRM2, we mutated F245 in RNP2 to a Y or an inosine (I). Both mutant proteins are expressed at wt levels *in vivo* (Supplementary Figure S1D) and cause a temperature sensitive growth phenotype (Supplementary Figure S1E). As *npl3-F245I* cells grow a bit better at 30°C than *npl3-F245Y* cells (Supplementary Figures S1E, S2E), we chose the former for further analysis.

### NMR spectroscopy and structural analysis

Structural modeling. We characterized the overall conformation and internal flexibility of the $Npl3^{120\text{-}280}$ protein using solution nuclear magnetic resonance (NMR) spectroscopy. The overall tumbling correlation time (Supplementary Figure S3C) derived from NMR $^{15}N$ relaxation experiments indicates that the two domains are arranged in a rather fixed orientation and do not reorient independently. The data also show that the N-terminal tail and the C-terminal tail regions flanking the two RRM domains are flexible (Supplementary Figure S3B-E). Interestingly, heteronuclear nuclear Overhauser effect (NOE) data (Supplementary Figure S3D) demonstrate that the linker connecting the two RRM domains has only reduced flexibility. The structural arrangement of the Npl3 tandem domains was then defined by inter-domain distance restraints derived from paramagnetic relaxation enhancement (PRE) experiments following our previously reported approach in a semi-rigid body structure calculation (see methods) (1) (Figure 3C, Supplementary Figure S5A-E). The calculated tandem RRM structure of Npl3 is in good agreement with experimental PRE and small angle X-ray scattering (SAXS) data (Supplementary Figure S5D). The structural model (Figure 3C) shows that the RNA-binding surfaces of the two RRM domains face towards each other, forming a positively charged surface (Figure 3D).

Npl3 RRM1 and RRM2 binding preference. We assessed the RNA-binding specificity of Npl3 RRM1 and RRM2 using NMR titrations with short single-stranded DNA oligos. It has been shown before that ssDNA ligands can well represent the RNA recognition by RRMs (2), especially in the case of SRSF1. The maximum NMR chemical shift perturbation observed with the RNP1 and RNP2 binding sites for RRM1 and within the non-canonical conserved motif in RRM2 is plotted in Figure 3B and Supplementary Figure S4A-F, for the oligo sequences indicated.

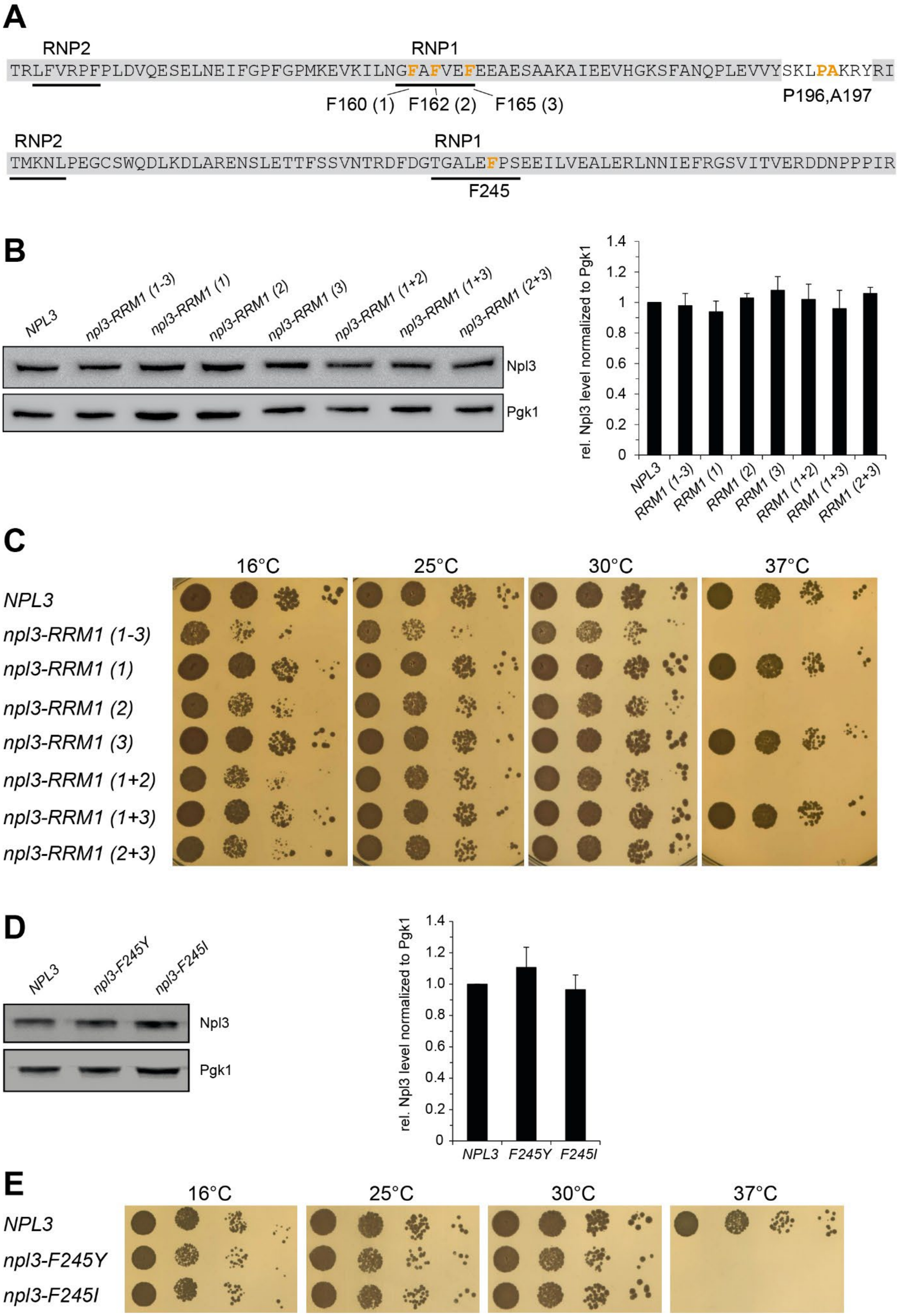
A
RNP2
RNP1
TRLFVRPFPLDVQESELNEIFGPFGPMKEVKILNGFAFVEFEEAESAAKAIEEVHGKSFANQPLEVVYSKLPAKRYRI
F160 (1)
F162 (2)
F165 (3)
P196,A197
RNP2
RNP1
TMKNLPEGCSWQDLKDLARENSLETTFSSVNTRDFDGTGALEFPSEEILVEALERLNNIEFRGSVITVERDDNPPPIR
F245
B
NPL3
npl3-RRM1 (1-3)
npl3-RRM1 (1)
npl3-RRM1 (2)
npl3-RRM1 (3)
npl3-RRM1 (1+2)
npl3-RRM1 (1+3)
npl3-RRM1 (2+3)
Npl3
Pgk1
rel. Npl3 level normalized to Pgk1
1.4
1.2
1
0.8
0.6
0.4
0.2
0
NPL3
RRM1 (1-3)
RRM1 (1)
RRM1 (2)
RRM1 (3)
RRM1 (1+2)
RRM1 (1+3)
RRM1 (2+3)
C
16°C
25°C
30°C
37°C
NPL3
npl3-RRM1 (1-3)
npl3-RRM1 (1)
npl3-RRM1 (2)
npl3-RRM1 (3)
npl3-RRM1 (1+2)
npl3-RRM1 (1+3)
npl3-RRM1 (2+3)
D
NPL3
npl3-F245Y
npl3-F245I
Npl3
Pgk1
rel. Npl3 level normalized to Pgk1
1.4
1.2
1.0
0.8
0.6
0.4
0.2
0
NPL3
F245Y
F245I
E
16°C
25°C
30°C
37°C
NPL3
npl3-F245Y
npl3-F245I

**Supplementary Figure S1.** Analysis of mutations in the RRM1 and RRM2 domains of Npl3. (**A**) Scheme of the RRM1-Linker-RRM2 region of Npl3 indicating the RNA-cross-linked amino acids of and mutations in Npl3. For the *npl3-RRM1* mutant F162 was mutated to tyrosine (Y), for *npl3-Linker* P196 and A197 were mutated to aspartic acid (D) and for *npl3-RRM2* F245 was mutated to isoleucine (I).

(**B**) Npl3 variants with mutations in the RRM1 domain are expressed at wt levels *in vivo*. Total Npl3 protein levels were determined by Western blot of whole-cell extracts from *NPL3* cells and cells expressing versions of Npl3 with the first (1), second (2) or third (3) phenylalanine mutated or combinations thereof. Left panel: Western blots, right panel: Quantification of three independent experiments. (**C**) Mutation of F162 to Y in the RRM1 of Npl3 is sufficient for the observed growth phenotype. F160, F162 and/or F165 (3) were mutated to Y either all three simultaneously (1-3), combinations of two mutations (1+2, 1+3 and 2+3) or the separately (1, 2 and 3). Mutation of F162 (2) confers the temperature sensitive phenotype.10-fold serial dilutions of wt cells and the indicated *npl3-RRM1* mutants were spotted onto YPD plates and grown for 2-3 days at the indicated temperatures. (**D**) Npl3 variant with mutations in the RRM2 domain of Npl3 are expressed at wt levels *in vivo*. Determination of total protein levels of npl3-F245Y and F245I mutants as in B. (**E**) Mutation of F245 to Y or I in the RRM2 of Npl3 causes a growth phenotype. Experiment as in C.

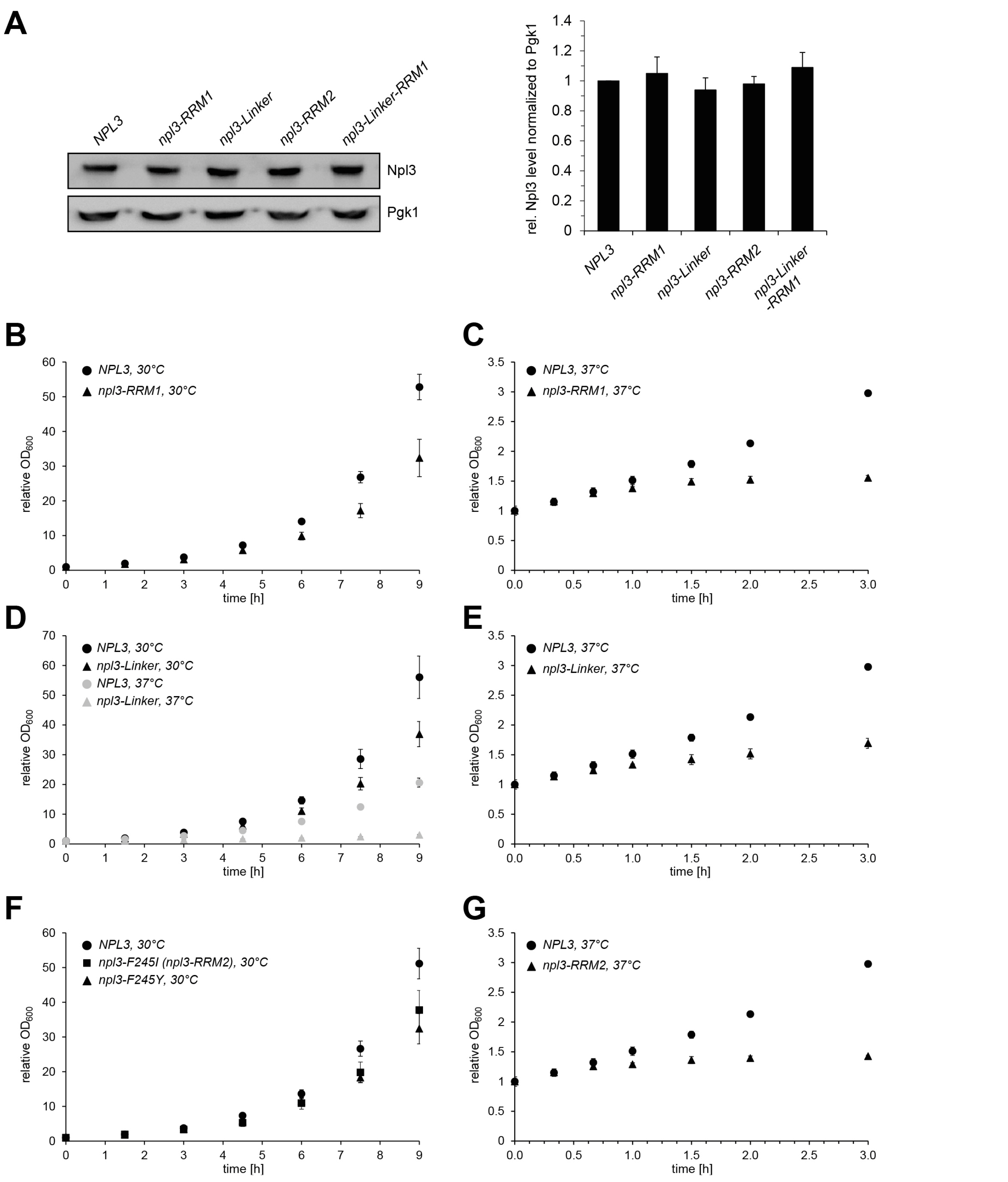


**Supplementary Figure S2.** Expression levels and growth curves of the three *npl3* mutants *npl3-RRM1*, *-Linker* and *-RRM2*. (**A**) Npl3 variant with mutations in the RRM1-, linker- and RRM2-domains of Npl3 are expressed at wt levels *in vivo*. Experiment as in Supplementary Figure S1B. (**B - G**) Growth curves of the *npl3-RRM1*, *-Linker* and *-RRM2* mutant at 30°C and 37°C as indicated.

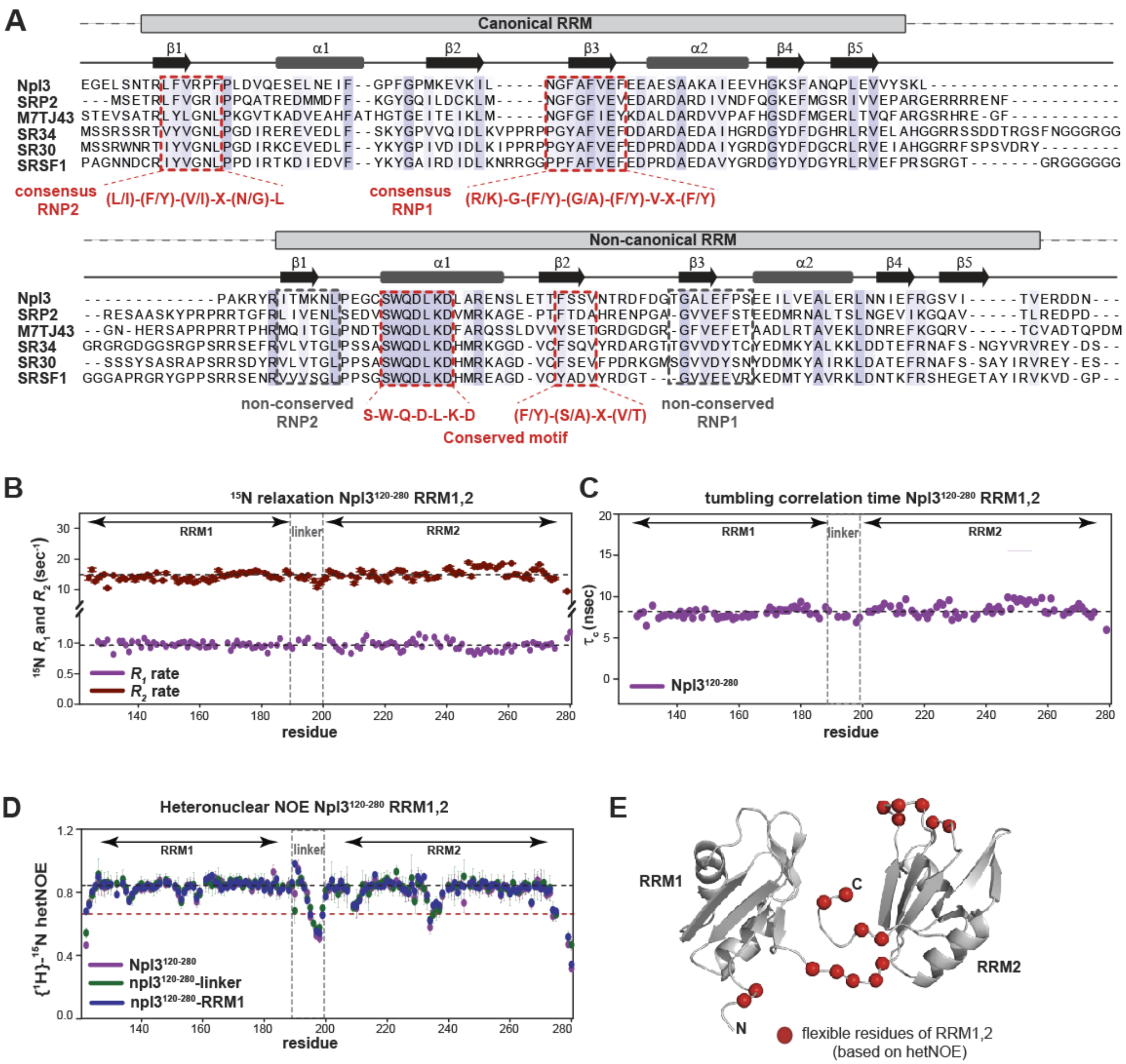


**Supplementary Figure S3.** Sequence conservation and NMR characterization of Npl3 RRM1,2. (**A**) Multiple sequence alignment of RRM domains from SR-like proteins across the various species using Clustal omega (https://www.ebi.ac.uk/Tools/msa/clustalo/) and Jalview (https://www.jalview.org/) tools. The red dashed line highlights the alignment of consensus RNP motifs in RRM1 and the non-canonical binding region in RRM2. (**B**) NMR $^{15}$N longitudinal ($R_1$) and transverse ($R_2$) relaxation rates at 900 MHz proton Larmor frequency. Average $R_1$ and $R_2$ rates are 0.97 sec$^{-1}$ and 14.8 sec$^{-1}$, respectively. (**C**) Residue-specific correlation times ($\tau_c$) calculated from $R_1$ and $R_2$ rates. The average correlation time is 8.1 ns. (**D**) Comparison of {$^1$H}-$^{15}$N heteronuclear NOE of wildtype and mutants of RRM1,2. Linker between RRM domains shown with gray box. (**E**) Residues with fast scale dynamics are highlighted on structure with red spheres.

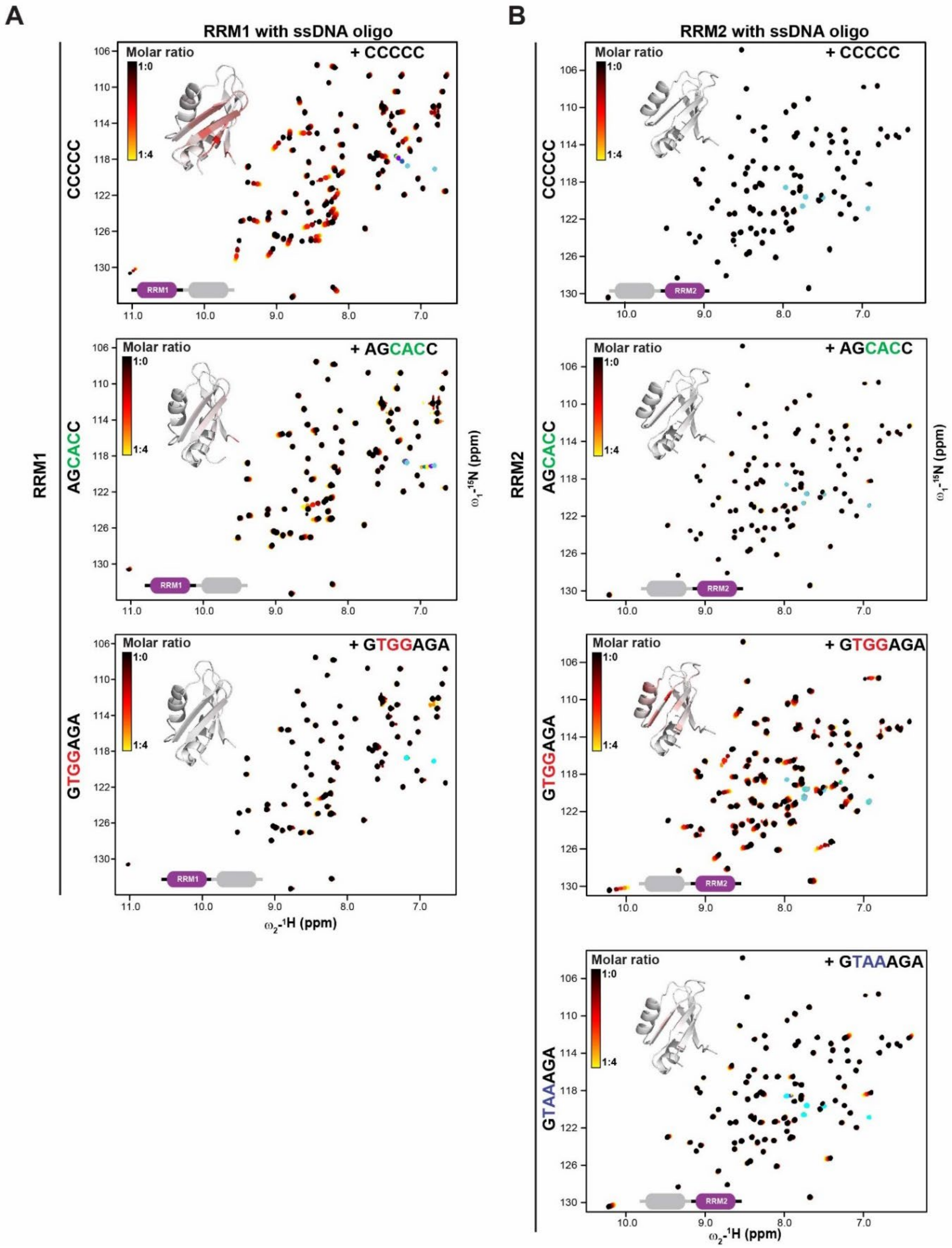
A
RRM1 with ssDNA oligo
B
RRM2 with ssDNA oligo
RRM1
RRM2
CCCCC
AGCACC
GTGGAGA
GTAAAGA
Molar ratio
1:0
1:4
+ CCCCC
+ AGCACC
+ GTGGAGA
+ GTAAAGA
RRM1
RRM2
ω1-15N (ppm)
ω2-1H (ppm)

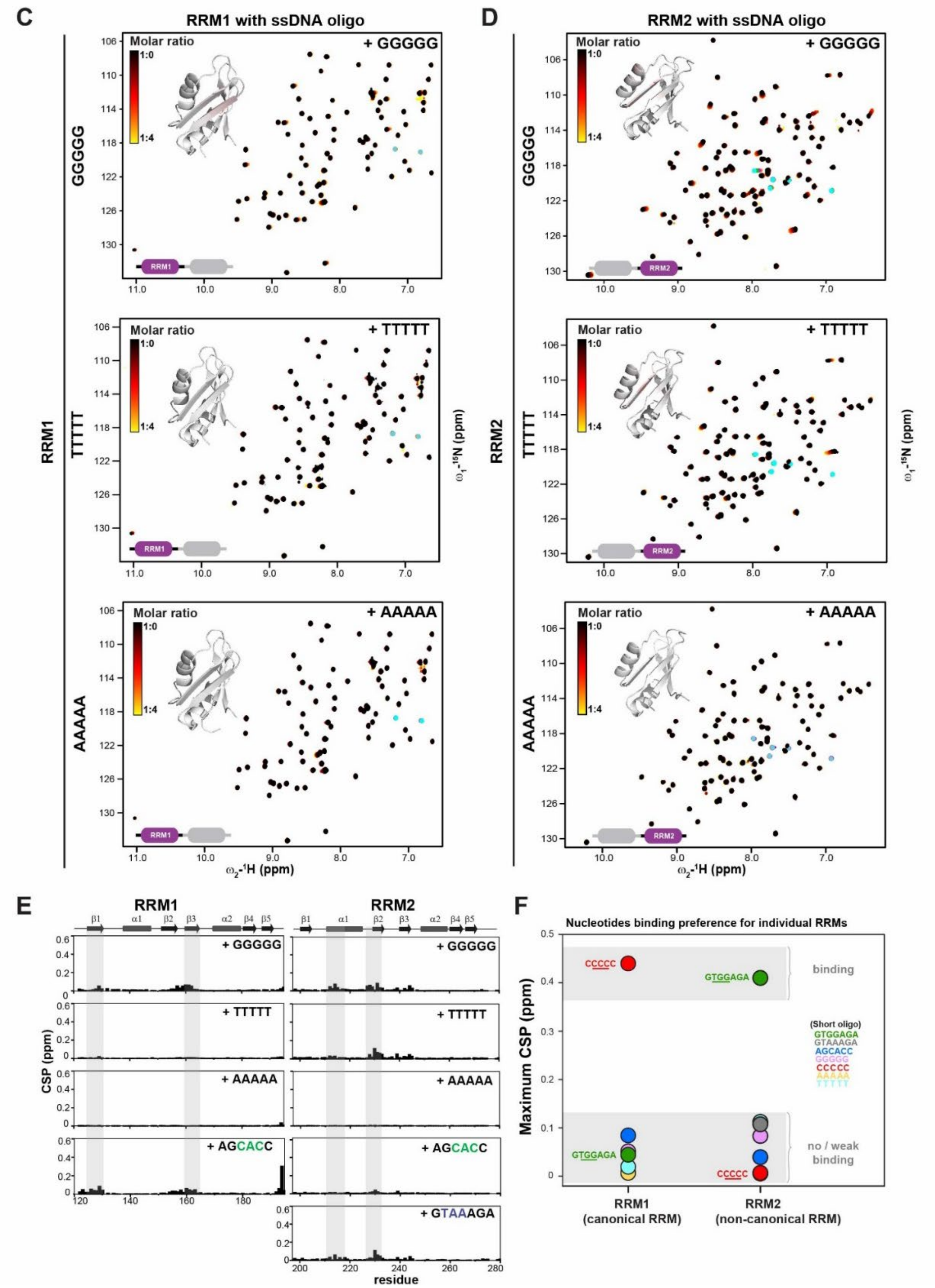


**Supplementary Figure S4.** NMR titrations to assess the RNA-binding preference of Npl3. Superposition of NMR $^{1}H$-$^{15}N$ correlation spectra of (**A,C**) RRM1 (left) and (**B,D**) RRM2 (right) titrated with various single-stranded DNA oligonucleotides. Spectra are colored black (free) and red to yellow (with increasing ligand concentration). Chemical shift perturbations (CSP) are mapped (red) onto the structure of the RRM domains. (**E**) Summary of the CSPs. RNP1, RNP2 and non-canonical conserved sites are highlighted with gray boxes. (**F**) Binding preferences for RRM1 (left) and RRM2 (right) based on maximum CSPs.

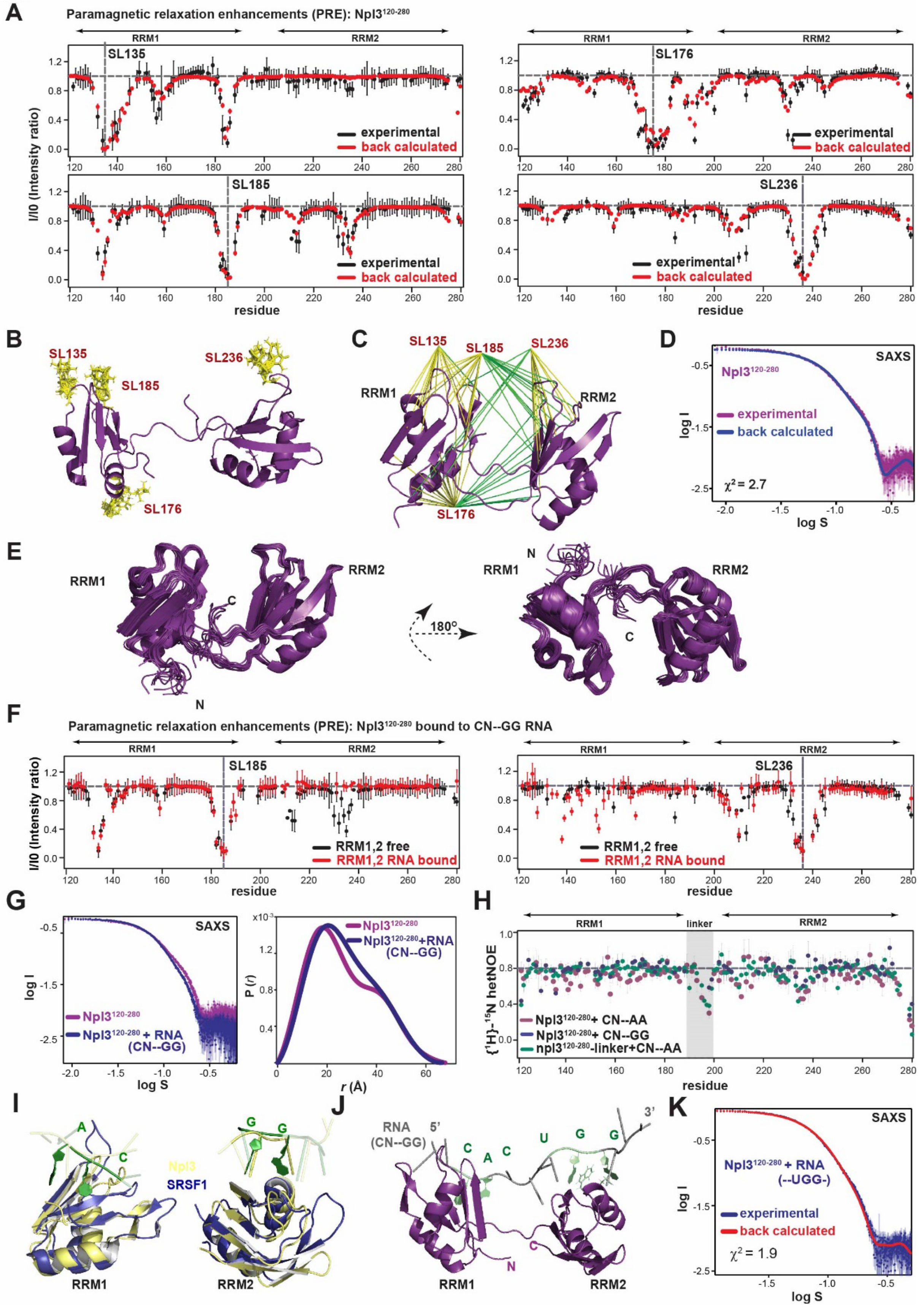

A
Paramagnetic relaxation enhancements (PRE): Npl3120-280
RRM1
RRM2
SL135
SL176
SL185
SL236
I/I0 (Intensity ratio)
experimental
back calculated
residue
B
SL135
SL185
SL236
SL176
C
SL135
SL185
SL236
RRM1
RRM2
SL176
D
SAXS
Npl3120-280
experimental
back calculated
χ2 = 2.7
log I
log S
E
RRM1
RRM2
N
C
180°
F
Paramagnetic relaxation enhancements (PRE): Npl3120-280 bound to CN--GG RNA
RRM1,2 free
RRM1,2 RNA bound
G
SAXS
Npl3120-280
Npl3120-280 + RNA (CN--GG)
P (r)
r (Å)
H
linker
{1H}-15N hetNOE
Npl3120-280+ CN--AA
Npl3120-280+ CN--GG
npl3120-280-linker+CN--AA
I
Npl3
SRSF1
RRM1
RRM2
J
RNA (CN--GG)
5'
3'
C A C U G G
N
C
K
SAXS
Npl3120-280 + RNA (--UGG-)
experimental
back calculated
χ2 = 1.9

**Supplementary Figure S5.** Structure of the Npl3 RRM domains free and RNA-bound. (**A**) Paramagnetic relaxation enhancements (PREs) from amide signal intensities in the oxidized and reduced state of spin-labeled protein (black) *vs.* PREs back-calculated from the final model (red). Four positions were spin-labeled individually by introducing mono-Cys variants for residue 135, 176,185 and 236. (**B**) Starting template for structure calculation with an ensemble of four copies for each spin label site indicated by yellow sticks. (**C**) Structural model of Npl3 RRM1,2 based on intra- (yellow lines) and inter-RRM (green lines) distance restraints derived from the experimental PRE data. (**D**) Comparison of experimental small angle X-ray scattering (SAXS) data with those back-calculated from the final structure. (**E**) Superposition of the ensemble of 15 lowest energy structures for the tandem RRM1,2 in two different views. (**F**) PRE data of spin-labeled Npl3 RRM1,2 bound to CN--GG RNA (red) compared to PRE data of the free protein (black) for spin labels at position 185 (RRM1) and 236 (RRM2). (**G**) Experimental SAXS data of free and RNA-bound Npl3 RRM1,2. (**H**) $\{^{1}H\}$-$^{15}N$ heteronuclear NOE of wild-type and linker mutant of RRM1,2 bound to "CN—AA" and "CN—GG" RNA. (**I**) Overlay of the SRSF1 RRM/RNA complex structures (dark blue) with of Npl3 (pale-yellow). CN and GG motifs are colored in green. (**J**) NMR and SAXS-derived structural model of the Npl3 tandem RRM domains with the "CN--GG" RNA. (**K**) Experimental and back calculated SAXS data from RNA-bound model on $Npl3^{120\text{-}280}$ are shown. The agreement is indicated by a $\chi^2$ of 1.9.

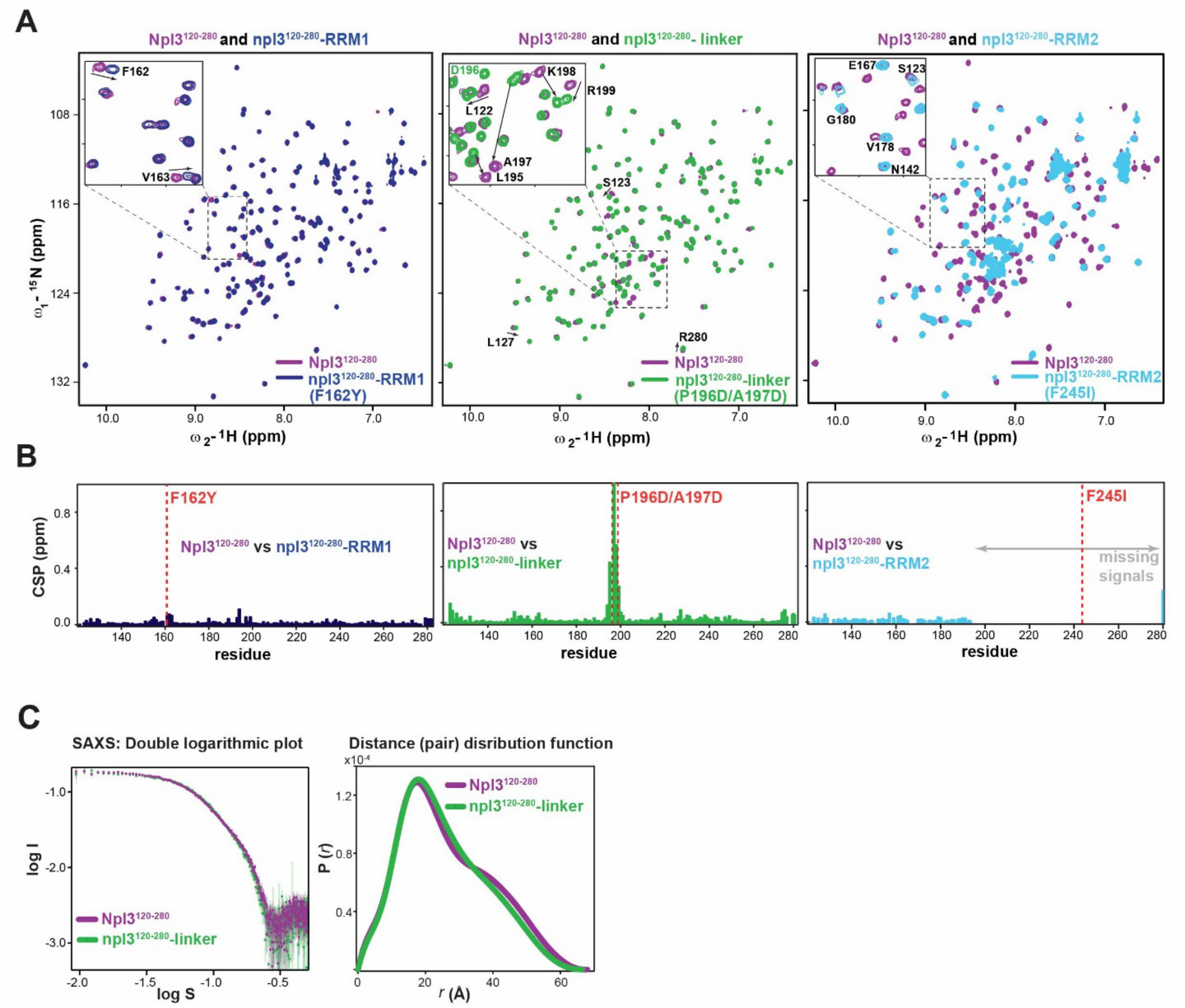


**Supplementary Figure S6.** Structural effects of mutations on Npl3$^{120\text{-}280}$ by NMR and SAXS. (**A**) NMR $^1$H-$^{15}$N correlation spectra (HSQC) of wildtype RRM1,2 (purple) superimposed with the RRM1 mutant (dark-blue), linker mutant (green) and RRM2 mutant (cyan), respectively. (**B**) Chemical shift perturbation (CSP) between spectra of wildtype Npl3$^{120\text{-}280}$ with npl1-RRM1, npl3$^{120\text{-}280}$-linker and npl3$^{120\text{-}280}$-RRM2 mutants, respectively. The mutated sites are indicated by red dashed lines. (**C**) SAXS data (left) and distance distribution function *P(r)* (right) for wild-type Npl3$^{120\text{-}280}$ and the linker mutant.

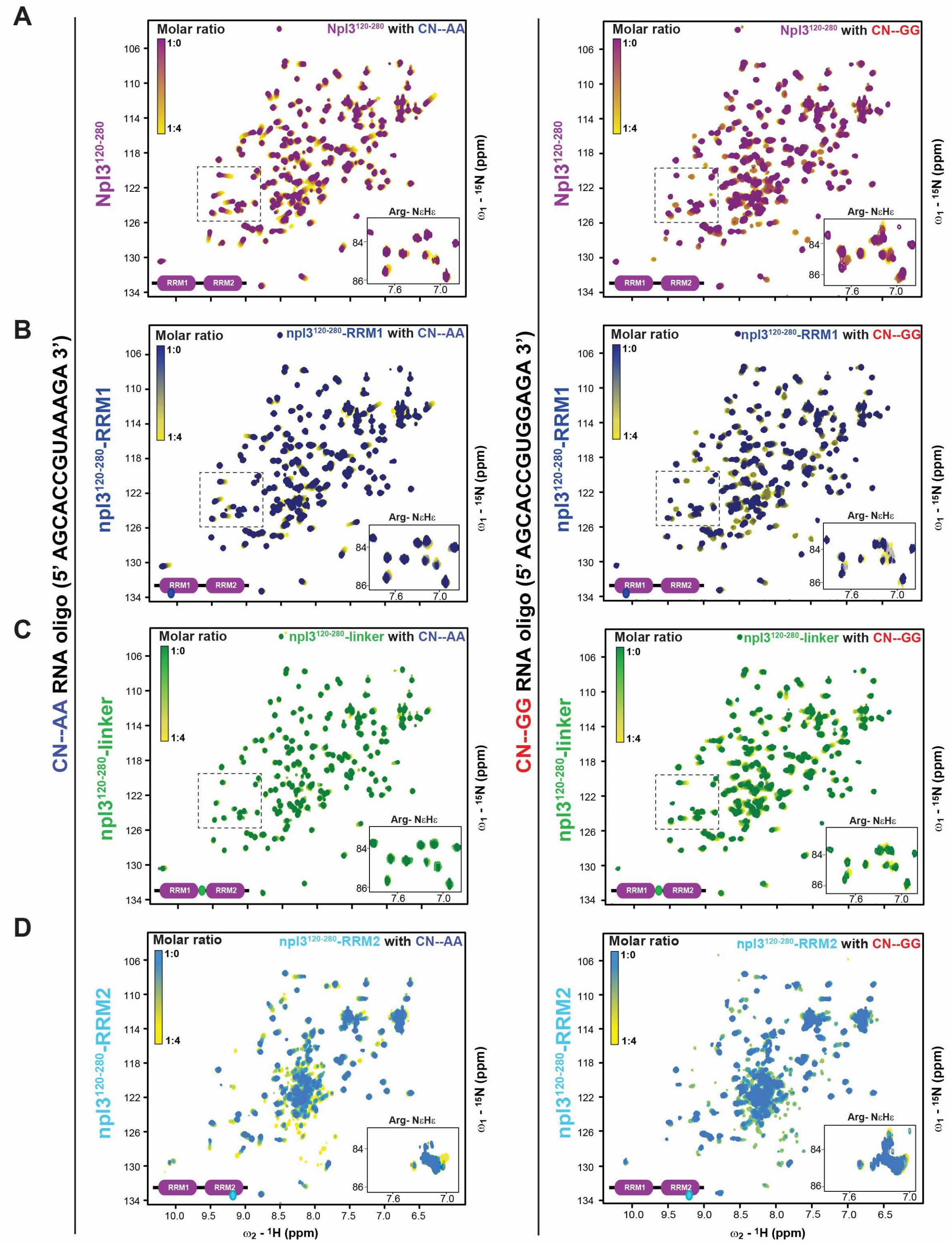


**Supplementary Figure S7.** RNA binding with wild-type and various mutants of Npl3$^{120-280}$ by NMR. Superposition of NMR $^{1}$H-$^{15}$N correlation spectra of (**A**) $^{15}$N labeled wildtype RRM1,2 (purple) (**B**) RRM1 (dark blue), (**C**) Linker –mutant (green) and (**D**) RRM2 (light blue) mutants titrated with "CN--AA" RNA (left) and the "CN--GG" RNA (right). Increasing chemical shift changes are colored from purple, green, dark or light blue (free form) to yellow (RNA-bound form). NMR signals of arginine NεHε side chains are shown as insets.

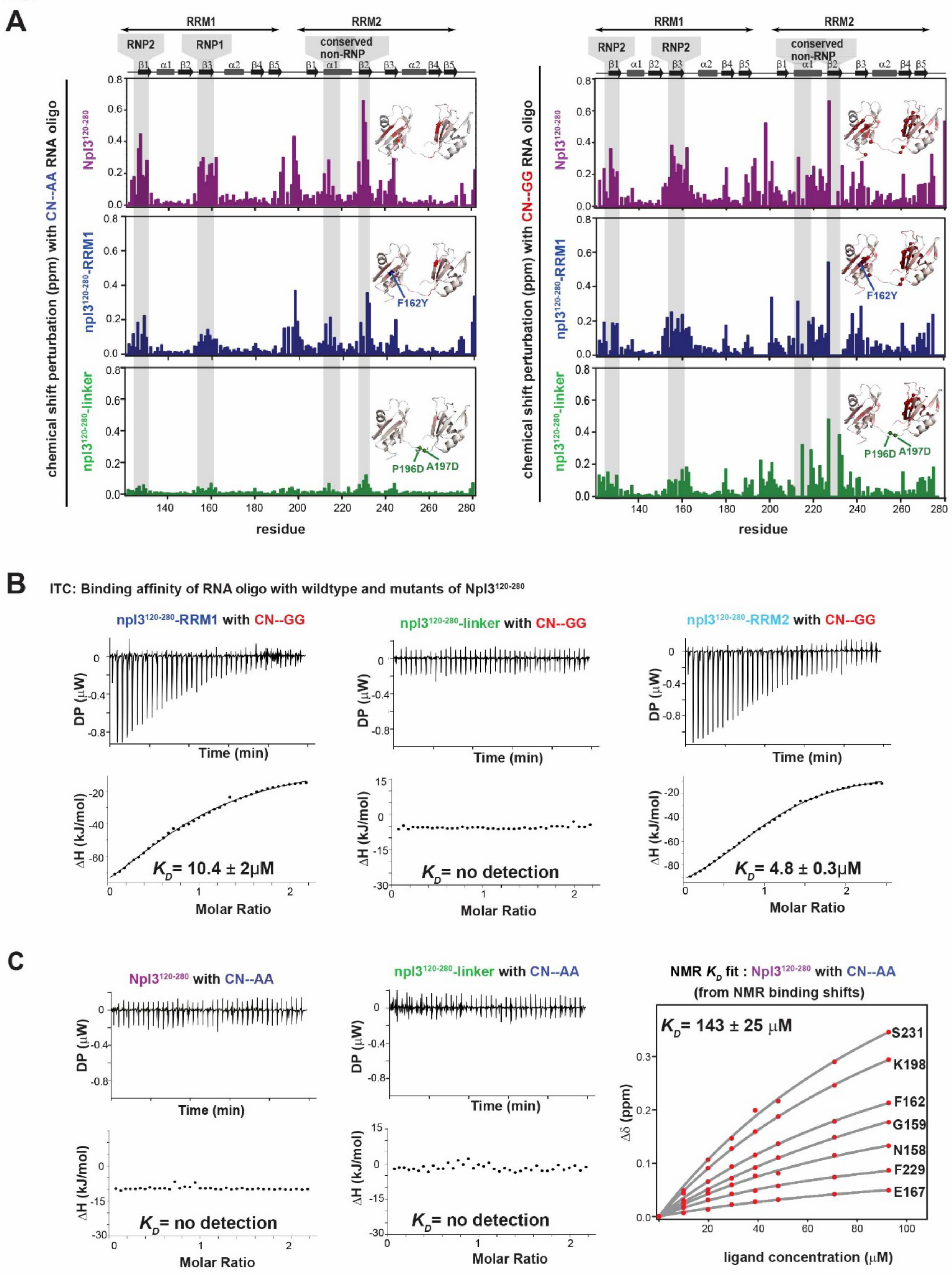

A
RRM1
RRM2
RNP2
RNP1
conserved non-RNP
chemical shift perturbation (ppm) with CN--AA RNA oligo
chemical shift perturbation (ppm) with CN--GG RNA oligo
Npl3120-280
npl3120-280-RRM1
npl3120-280-linker
F162Y
P196D
A197D
residue
B
ITC: Binding affinity of RNA oligo with wildtype and mutants of Npl3120-280
npl3120-280-RRM1 with CN--GG
npl3120-280-linker with CN--GG
npl3120-280-RRM2 with CN--GG
DP (μW)
Time (min)
ΔH (kJ/mol)
Molar Ratio
KD= 10.4 ± 2μM
KD= no detection
KD= 4.8 ± 0.3μM
C
Npl3120-280 with CN--AA
npl3120-280-linker with CN--AA
KD= no detection
KD= no detection
NMR KD fit : Npl3120-280 with CN--AA
(from NMR binding shifts)
KD= 143 ± 25 μM
S231
K198
F162
G159
N158
F229
E167
Δδ (ppm)
ligand concentration (μM)

**Supplementary Figure S8.** NMR and isothermal titration calorimetry (ITC) analysis of RNA binding to wild-type and mutant Npl3. (**A**) CSP plot for wildtype (purple), npl3-RRM1 (dark blue) and npl3-linker (green) with "CN--AA" (left) and "CN-GG" (right) RNAs from spectra shown in Supplementary Figure S6. Conserved sequence motifs (RNP1, RNP2 in RRM1 and the non-canonical site in RRM2) are highlighted with gray boxes. The inset shows mapping of CSPs (red) onto the tandem RRM1,2 structure. (**B**) ITC data shown for the binding of "CN--GG" RNA to Npl3$^{120\text{-}280}$-RRM1 (left), Npl3$^{120\text{-}280}$-linker (middle) and Npl3$^{120\text{-}280}$-RRM2 (right), respectively. (**C**) ITC data shown for binding of "CN--AA" RNA with Npl3$^{120\text{-}280}$ (left) and Npl3$^{120\text{-}280}$-linker (middle). $K_D$ calculated from NMR titrations (right panel) upon addition of "CN—AA" RNA to npl3$^{120\text{-}280}$ (see Supplementary Figure S6A).

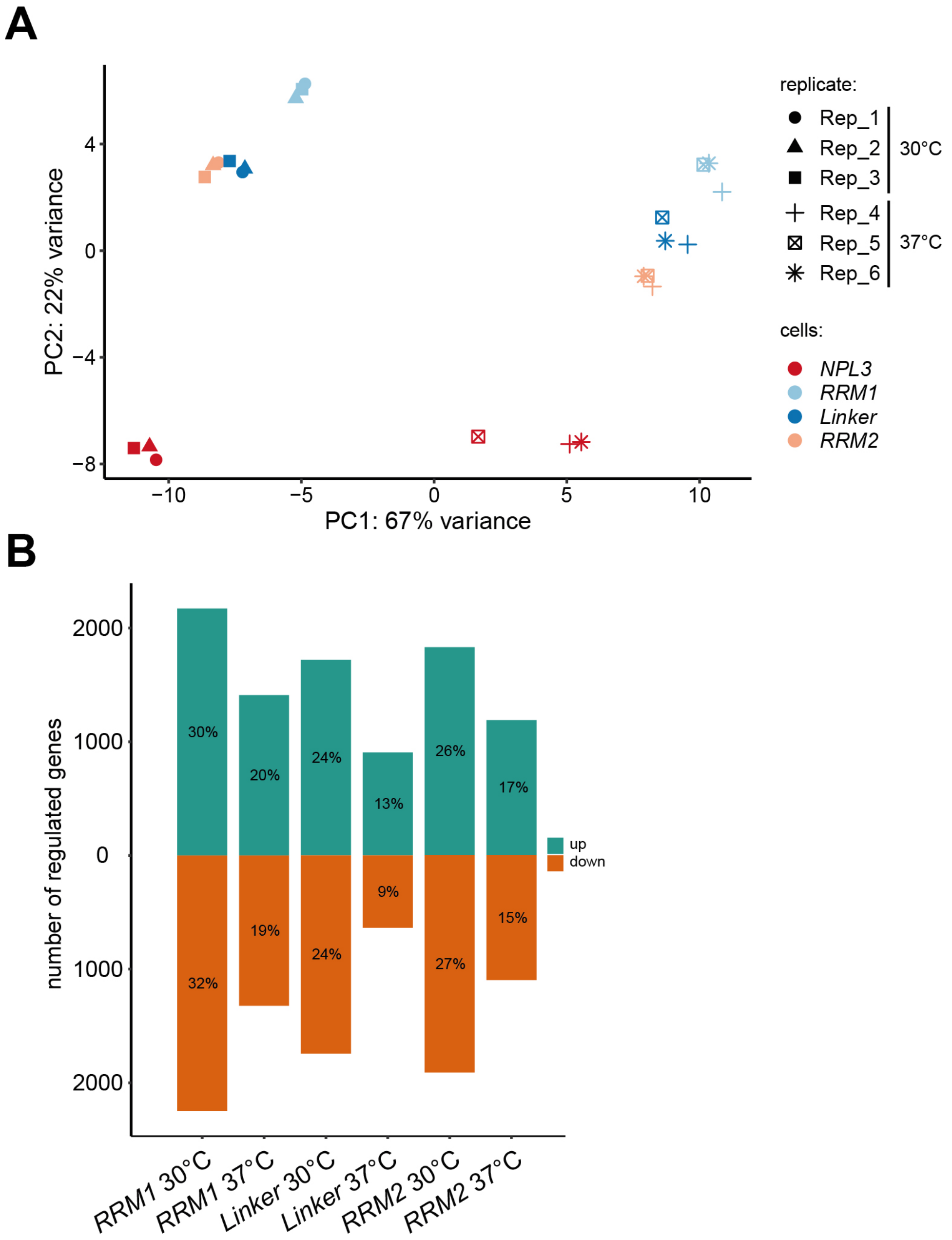


**Supplementary Figure S9.** Mutations in *NPL3* lead to substantial changes in transcript abundance. (**A**) Principle component analysis of regularized-logarithm transformed RNA-seq read counts based on 500 most variable genes across all samples. Scatter plot shows the first two principle components (PC1 and PC2) to evaluate the variance between wt and *npl3* mutants at 30°C and 37°C. Each condition comprises three biological replicates. (**B**) Hundreds of transcripts change in abundance in the *npl3* mutants. Bar diagram depicts number and percentages of significantly up-regulated (green) and down-regulated (orange) genes (adjusted p-value $< 0.05$) in the three *npl3* mutants at 30°C and 37°C.

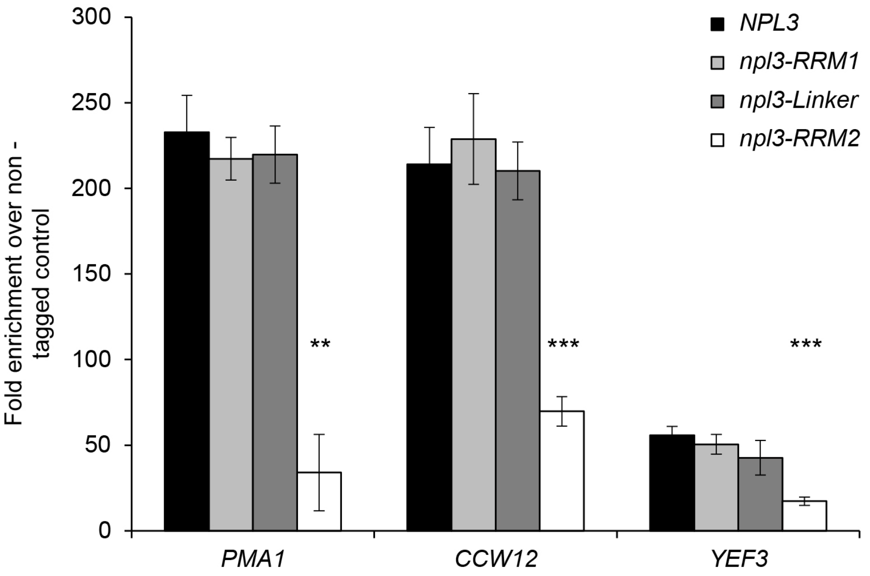


**Supplementary Figure S10.** RNA binding of the CBC subunit Cbc2 is not reduced in *npl3-RRM1* and *-Linker* cells *in vivo*. TAP-tagged Cbc2 was immunoprecipitated from whole-cell extracts of *NPL3*, *npl3-RRM1*, *-Linker* and *-RRM2* cells. The amount of co-immunoprecipitated RNA was quantified by RT-qPCR for the indicated transcripts. The amount of co-immunoprecipitated mRNA was calculated as the enrichment over the amount of RNA in a control experiment with a strain expressing non-tagged Cbc2.

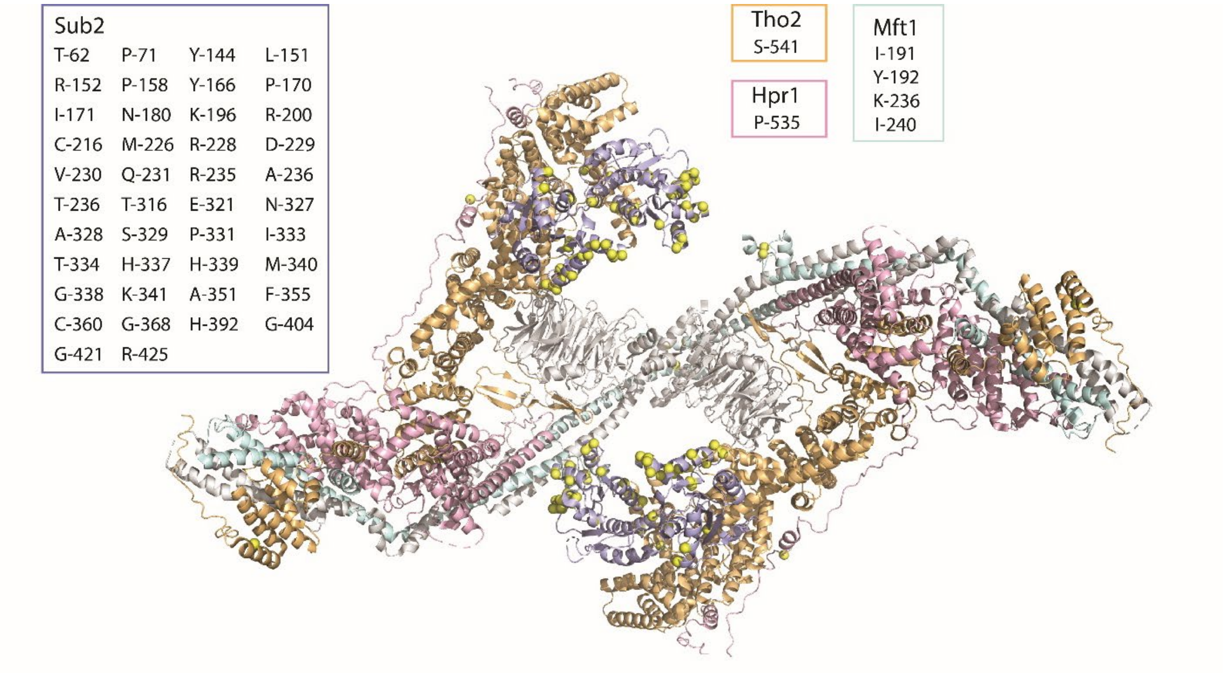


**Supplementary Figure S11.** 3D structure of THO-Sub2 complex (3) with RNA crosslinking sites in Tho2, Hpr1, Mft1 and Sub2 (Supplementary Table S1 and S2). Amino acids crosslinked to RNA are depicted as yellow spheres. Coloring scheme of the proteins with crosslinking sites is illustrated. The crosslinking sites of the respective proteins are listed within the inlets.

**Supplementary Table S1. Amino Acids identified to be cross-linked to RNA *in vivo* by RNP$^{XL}$**
Protein names with asterisk indicate those peptides with their crosslinked amino acids that have been previously identified (4). Asterisks in parentheses mark the crosslinked amino acids identified in this study.

| Protein | Cross-linked peptide | Cross-linked amino acid | Source |
|---|---|---|---|
| Npl3 | 127-LFVRPFPLDVQESELNEIFGPFGPMK-152 | 127-152 | Cbc2 |
| Npl3* | 156-ILNGFAFVEFEEAESAAK-173 | F160 or F162 or F165 | Cbc2 |
| Npl3* | 182-SFANQPLEVVYSKLP(*)AK-198 | P196 | Hpr1, Sub2, Cbc2, Npl3, Mtr2, oligo(dT) |
| Npl3* | 222-ENSLETTFSSVNTR-235 | F229 or S230 and R235 | Cbc2, Hpr1, Mtr2, Npl3 |
| Npl3 | 273-DDNPPP(*)IR-280 | P278 | Cbc2, Npl3 |
| Npl3 | 273-DDNPPPIR(*)R-281 | R280 | Sub2, Hpr1, Tho1, Cbc2, Mtr2, Nab2, Npl3 |
| Npl3 | 371-NDYGP(*)PR-377 | P375 | Hpr1, Cbc2 |
| Npl3 | 392-GDYG(*)P(*)PR-398 | G395 and P396 | Cbc2, Hpr1 |
| Npl3 | 385-GGYDGPRGDYGPPRDAYR(*)TR-404 | R402 | Sub2 |
| Sto1 | 7-RGDF(*)DEDENYR-17 | R7 | Cbc2 |
| Sto1 | 18-DF(*)RPR-22 | F19 | Cbc2 |
| Sto1 | 18-DFRP(*)R-22 | P21 | Cbc2 |
| Sto1 | 107-NNVAGKSIINYFFEELQK-124 | K112 or S113 | Thp1 |
| Sto1 | 578-K(*)NDLYFR-584 | K578 | Cbc2 |
| Cbc2 | 17-RL(*)DTPSR-23 | L18 | Cbc2 |
| Cbc2 | 31-R(*)NPNGLQELR-40 | R31 | Sub2, Cbc2 |
| Cbc2 | 134-GK(*)SGGQVSDELR-145 | K135 | Cbc2 |
| Cbc2 | 146-FDFDASRGGFAIPFAER-162 | 146-162 | Cbc2 |
| Cbc2 | 163-VGVP(*)HSR-169 | P166 | Cbc2 |
| Tho2 | 1183-LPSSALIGH(*)LK-1193 | H1191 | Hpr1 |
| Tho2 | 1357-TLIQNPQNP(*)DFAEK-1370 | P1365 | Hpr1, Sub2 |
| Hpr1 | 534-IP(*)TGLDK-540 | P535 | Thp1 |
| Mft1 | 189-YRIYDDFSK-197 | I191 or Y192 | Hpr1, Sub2 |
| Gbp2 | 98-GR(*)TLGPIVER-107 | R-99 and G-98 or G-102 | Hpr1, Sub2 |
| Gbp2 | 100-TLGP(*)IVER-107 | P-103 | Sub2 |
| Hrb1 | 51-FADTY(*)RGSR-59 | Y55 | Hpr1 |
| Hrb1 | 137-GDYGP(*)LLAR-145 | P141 | Hpr1, Sub2 |
| Hrb1 | 190-ADIITSRGHHR-200 | S195 or R196 | Hpr1, Cbc2, Mtr2 |
| Sub2 | 53-K(*)GSYVGIHSTGFK-65 | K53 | Sub2 |
| Sub2 | 53-KG(*)SY(*)VGIHSTGFK-65 | G54 and Y56 | Sub2 |
| Sub2 | 54-GSYVGIHST(*)GFKDFLLKPELSR-75 | T62 | Sub2 |
| Sub2 | 66-DFLLKP(*)ELSR-75 | K70 | Sub2 |
| Sub2 | 141-ELAY(*)QIR-147 | Y144 | Sub2 |
| Sub2* | 162-TAVFYGGTP(*)ISK-173 | P170 | Sub2, Hpr1, oligo(dT) |

| | | | |
|---|---|---|---|
| Sub2 | 180-NKDTAPHIVVATPGR-194 | N180 or K181 | Sub2 |
| Sub2 | 195-LK(*)ALVR-200 | K196 | Sub2 |
| Sub2 | 197-ALVR(*)EK-202 | R200 | Hpr1, Mtr2 |
| Sub2 | 201-EKYIDLSHVK-210 | E201 or K202 | Sub2 |
| Sub2 | 203-Y(*)IDLSHVK-210 | Y203 | Sub2 |
| Sub2 | 211-NFVIDEC(*)DK-219 | C217 | Sub2 |
| Sub2 | 228-RDVQEIFR-235 | R228 or D229 | Sub2 |
| Sub2 | 228-RDVQEIFR(*)ATPR-239 | R235 | Sub2 |
| Sub2 | 260-RF(*)LQNPLEIFVDDEAK-275 | F261 | Sub2 |
| Sub2 | 315-STTRANELTK-324 | T316 or T317 | Hpr1, Mtr2 |
| Sub2* | 325-LLNASNFPAITVHGH(*)MK-341 | H339 | Sub2, oligo(dT), Hpr1 |
| Sub2 | 351-AFKDF(*)EK-357 | F355 | Sub2 |
| Sub2 | 359-IC(*)VSTDVFGR-368 | C360 | Sub2 |
| Sub2 | 369-G(*)IDIER-374 | G369 | Sub2 |
| Sub2 | 375-INLAINYDLTNEADQYLH(*)R-393 | H392 | Sub2 |
| Sub2 | 404-G(*)LAISFVSSK-413 | G404 | Sub2 |
| Sub2 | 414-EDEEVLAK(*)IQER-425 | K421 | Sub2 |
| Yra1 | 8-SLDEIIG(*)SNKAGSNR-22 | G14 and K17 or A18 | Sub2 |
| Yra1 | 8-SLDEIIGSNKAG(*)SNR-22 | G19 | Sub2, Hpr1, Mtr2 |
| Yra1 | 25-V(*)G(*)G(*)T(*)R(*)GNGPR-34 | 25-29 | Mtr2, Hpr1, Tho1, Cbc2 |
| Yra1 | 36-VG(*)K(*)QVGSQR-44 | G37 and K38 | Cbc2, Mtr2, Hpr1, Nab2 |
| Yra1 | 45-R(*)S(*)LP(*)NR-50 | R45, S46 and P48 | Hpr1, Tho1, Cbc2, Mtr2, Nab2 |
| Yra1 | 51-R(*)GP(*)IR-55 | R51 and P53 | Sub2, Hpr1, Tho1, Mtr2, Cbc2, Nab2, Npl3 |
| Yra1 | 57-NTR(*)APPNAVAR-67 | R59 | Sub2, Tho1, Cbc2, Mtr2 |
| Yra1* | 57-NTRAP(*)PNAVAR-67 | P61 | oligo(dT), Tho1, Hpr1, Mtr2, Nab2, Npl3 |
| Yra1 | 57-NTRAPP(*)NAVAR-67 | P62 | Hpr1, Cbc2, Mtr2, Nab2 |
| Yra1 | 60-APPNAVAR(*)VAK-70 | R67 | Sub2, Nab2, Mtr2 |
| Yra1 | 68-VAK(*)LLDTTR-76 | K70 | Hpr1, Sub2, Mtr2 |
| Yra1 | 80-VNVEGLPRDIK-90 | P86 or R87 | Hpr1, Sub2 |
| Yra1 | 108-VLLSYNER(*)GQSTGMANITFK-127 | R115 | Sub2 |
| Yra1 | 139-FNGSPIDGGRSR-150 | 146-GGRSR-150 | Sub2, Hpr1, Mtr2, Nab2, Npl3 |
| Yra1* | 153-LNLIVDP(*)NQR(*)PVK-165 | P159 and R162 | Hpr1, Sub2, oligo(dT) |
| Yra1* | 153-LNLIVDPNQRP(*)VK-165 | P163 | Sub2, Mtr2, oligo(dT) |
| Yra1 | 178-GGNAP(*)R(*)P(*)VK-186 | P182, R183 and P184 | Hpr1, Tho1, Cbc2, Nab2, Mtr2 |
| Yra1 | 210-KSLEDLDK-217 | K210 or S211 | Sub2 |
| Yra2 | 38-LGFAPSDAASR(*)SK-50 | R48 | Cbc2 |
| Tho1 | 75-KEVSSEPK-82 | 75-82 | Nab2 |
| Tho1 | 129-ALDLLNK(*)K-136 | K135 | Tho1 |
| Tho1 | 140-ANKF(*)GQDQADIDSLQR-155 | F143 | Tho1 |
| Tho1 | 181-KNEPESGNNGKFK-193 | G190 or K191 or F192 | Tho1 |

| | | | |
|---|---|---|---|
| Tho1 | 214-SGYRR-218 | G215 or Y216 or R217 | Tho1 |
| Nab2 | 210-GGGAVGK(*)NR-218 | K216 | Nab2 |
| Nab2 | 314-TREEFQK-320 | 314-320 | Nab2 |
| Nab2 | 322-KADL(*)LAAK-329 | L325 | Nab2 |
| Nab2 | 322-KADLLAAK(*)R-330 | K329 | Nab2 |
| Nab2 | 394-EVKPISQK(*)K-402 | K401 | Nab2 |
| Nab2 | 403-AAPPP(*)VEK-410 | P407 | Nab2 |
| Nab2 | 411-SLEQC(*)K-416 | C415 | Nab2 |
| Nab2 | 417-FGTHC(*)TNK-424 | C421 | Nab2 |
| Nab2 | 433-SHIMC(*)R-438 | C437 | Nab2 |
| Sac3 | 460-ALSH(*)TLNK-467 | H463 | Thp1 |
| Sac3 | 527-TYLTC(*)LER-534 | C531 | Thp1 |
| Sac3 | 862-NLIFSPVNDEFNK(*)FATHLTK-881 | K874 | Thp1 |
| Sac3 | 1084-YDK(*)TLR-1089 | K1086 | Thp1 |
| Sac3 | 1116-KMLEKEK(*)-1122 | K1122 | Nab2 |
| Cdc31 | 51-ALGFELP(*)KR-59 | P57 | Thp1 |
| Cdc31 | 113-ISIK(*)NLR-119 | K116 | Thp1 |
| Cdc31 | 121-VAK(*)ELGETLTDEELR-135 | K123 | Thp1 |
| Mex67 | 428-YNH(*)GYNSTSNNK-439 | H430 | Mtr2 |
| Mtr2 | 102-VR(*)FDESGR-109 | R-103 | Mtr2 |
| Mtr2 | 38-LAQFVQLFNPNNCR-51 | C-50 | Mtr2 |
| Pab1* | 307-QYEAYRLEK-315 | Y308 and Y311 | Cbc2, Npl3, Nab2 |
| Cbf5 | 467-KSEDGDSEEKK-477 | S-468 or S-473 | Cbc2 |
| Cbf5 | 420-EVETE(*)KEEVK-42 | E-424 | Sub2 |
| Msl5 | 90-KNRSPSPPPVY(*)DAQGK-105 | Y-100 | Tho1 |
| Msl5 | 268-E(*)DNRPCPICGLKDHK-282 | E-268 | Sub2 |
| Nam8 | 237-VGPTSGQQQH(*)VSGNNDYNR-25 | H-246 and Q-245 or Q-244 or Q-243 | Cbc2 |
| Prp19 | 140-SSQQAV(*)AITR-149 | V-145 | Mtr2 |
| Yhc1 | 44-DIINKHNHK-52 | N-47 or N-50 | Cbc22 |
| Yhc1 | 55-HIGKRGR-61 | H-55 | Hpr1 |

**Supplementary Table S2. Identified crosslinks of TREX components**
Index numbers indicate the MS2 spectrum identified in the respective idXml file of the affinity purifications. See Excel table.

**Supplementary Table S3. Crosslinks identified by a database search against the entire yeast database**
Index numbers list the MS2 spectra of crosslinks in the respective Affinity purification with their respective idXml files. See Excel table.

**Supplementary Table S4A. List of oligonucleotides used NMR binding studies**

| | Npl3 domains | oligo | NMR CSP[b] | Binding[c] |
|---|---|---|---|---|
| DNA oligonucleotides (single stranded)[a] | RRM1 | TTTTT | 0.01 | no |
| | | GGGGG | 0.05 | no |
| | | AAA AA | 0.01 | no |
| | | CCCCC | 0.44 | yes |
| | | AGCCCC | 0.31 | yes |
| | | AGCACC | 0.08 | yes |
| | RRM2 | TTTTT | 0.11 | yes |
| | | CCCCC | 0.00 | no |
| | | AAA AA | 0.01 | no |
| | | GGGGG | 0.08 | yes |
| | | GGGGAGA | 0.05 | no |
| | | GTGGGGA | 0.06 | no |
| | | GTGGAGA | 0.41 | yes |
| | | GTAAAGA | 0.11 | yes |
| | RRM1,2 | AGCACCGTGGAGA | 0.34 | yes |

[a] DNA oligos were used as proxy for RNA

[b] Maximal NMR chemical shift perturbation observed at 4-fold excess of the oligonucleotide

[c] Above a CSP threshold ≤ 0.08

**Supplementary Table S4B. Isothermal titration calorimetry (ITC) and NMR binding assays for the protein-RNA interactions**

| Protein | RNA | $K_D$ (μM) | N | ΔH(kJ/mol) | TΔS (kJ/mol) | ΔG (kJ/mol) |
|---|---|---|---|---|---|---|
| $Npl3^{120-280}$ | CN--GG | 0.66 ± 0.04 | 0.9 | -67.5 ± 0.8 | -32.2 | -35.3 |
| $npl3^{120-280}$-Linker | CN--GG | Not detected | - | - | - | - |
| $npl3^{120-280}$-RRM1 | CN--GG | 10.4 ± 2 | 0.97 | -127 ± 15.6 | -98.2 | -28.5 |
| $npl3^{120-280}$-RRM2 | CN--GG | 4.8 ± 0.3 | 1.08 | -118 ± 3.2 | -87.8 | -30.4 |
| $Npl3^{120-280}$ | CN--AA | Not detected | - | - | - | - |
| $npl3^{120-280}$-Linker | CN--AA | Not detected | - | - | - | - |
| $Npl3^{120-280}$ | CN--AA | 143 ± 25 (by NMR) | - | - | - | - |

**Supplementary Table S5. Small angle X-ray scattering of free and RNA bound RRM1,2 of Npl3**

| Sample | $R_g$ (Angstrom) | $D_{max}$ (Angstrom) |
|---|---|---|
| $Npl3^{120-280}$ | 20.4 ± 0.12 | 67.7 |
| $Npl3^{120-280}$ + CN--GG | 20.4 ± 0.16 | 63.6 |

| | | |
|---|---|---|
| npl3$^{120-280}$-Linker | 19.0 ± 0.10 | 66.5 |

### Supplementary Table S6. Structural statistics

| Npl3$^{120-280}$ RRM1,2 free form | Statistics |
|---|---|
| Intra-domain PRE restraints | 69 |
| Inter-domain PRE restraints | 31 |
| Distance violations | 4 (<3 Å) |
| Average RMSD for 15 lowest energy structures[a] | 0.93 Å |
| PRE quality factor[b] | 0.10 |
| Agreement with SAXS data ($\chi^2$) | 2.70 |
| Backbone structural quality (Ramachandran plot) | 82.8% (allowed)<br>14.8% (additionally allowed),<br>1.6 % (generously allowed)<br>0.8% (dis-allowed) |

[a] out of 100 structures calculated
[b] see methods

### Supplementary Table S7. Differential expression analysis using DESeq2

See Excel table.

### Supplementary Table S8. Quantitative MS proteomic data of the Cbc2-TAP purifications from *npl3-Linker* and wt cells.

See Excel table.

### Supplementary Table S9. Yeast Strains

| Yeast Strain | Genotype | Reference |
|---|---|---|
| RS453 | MAT a, ade2-1, his3-11,15, ura3-52, leu2-3,112, trp1-1, can1-100, GAL+ | Sträßer K. 2000 |
| *SUB2-TAP* | MAT a, ade2-1, his3-11,15, ura3-52, leu2-3,112, trp1-1, can1-100, GAL+, SUB2-CBP-TEV-2x protA::TRP1-KL | Sträßer K. 2002 |
| *HPR1-FTpA* | MAT a, ade2-1, his3-11,15, ura3-52, leu2-3,112, trp1-1, can1-100, GAL+, HPR1-FLAG-TEV-2x protA::HIS3MX4 | This study |
| *TAP-NPL3* | MAT a, ade2-1, his3-11,15, ura3-52, leu2-3,112, trp1-1, can1-100, GAL+, 2x protA-TEV-NPL3::TRP1-KL | This study |
| *THO1-FTpA* | MAT a, ade2-1, his3-11,15, ura3-52, leu2-3,112, trp1-1, can1-100, GAL+, THO1-FLAG-TEV-2x protA::HIS3MX4 | This study |
| *NAB2-FTpA* | MAT a, ade2-1, his3-11,15, ura3-52, leu2-3,112, trp1-1, can1-100, GAL+, NAB2-FLAG-TEV-2x protA::HISMX4 | This study |
| *MTR2-FTpA* | MAT a, ade2-1, his3-11,15, ura3-52, leu2-3,112, trp1-1, can1-100, GAL+, MTR2-FLAG-TEV-2x protA::HIS3MX4 | This study |
| *CBC2-FTpA* | MAT a, ade2-1, his3-11,15, ura3-52, leu2-3,112, trp1-1, can1-100, GAL+, CBC2-FLAG-TEV-2x protA::HIS3MX4 | This study |
| *THP1-TAP* | MAT a, ade2-1, his3-11,15, ura3-52, leu2-3,112, trp1-1, can1-100, GAL+, THP1-CBP-TEV-2x protA::TRP1-KL | This study |
| W303 | MATa, ura3-1, trp1-1, his3-11,15, leu2-3,112, ade2-1, can1-100 GAL+ | Thomas B.J. 1989 |

| | | |
|---|---|---|
| *npl3* shuffle | MATa, ura3-1, trp1-1, his3-11,15, leu2-3,112, ade2-1, can1-100 GAL+, NPL3::HIS3, pRS316-NPL3 | This study |
| *npl3* shuffle *CBC2-TAP* | MATa, ura3-1, trp1-1, his3-11,15, leu2-3,112, ade2-1, can1-100 GAL+, NPL3::HIS3, pRS316-NPL3, CBC2-CBP-TEV-2x protA::TRP1-KL | This study |
| *npl3* shuffle *CBC2-TAP* *MEX67-HA* | MATa, ura3-1, trp1-1, his3-11,15, leu2-3,112, ade2-1, can1-100 GAL+, NPL3::HIS3, pRS316-NPL3, CBC2-CBP-TEV-2x protA::TRP1-KL, MEX67-HA::KanMX4 | This study |
| *npl3* shuffle *CBC2-TAP* *HPR1-HA* | MATa, ura3-1, trp1-1, his3-11,15, leu2-3,112, ade2-1, can1-100 GAL+, NPL3::HIS3, pRS316-NPL3, CBC2-CBP-TEV-2x protA::TRP1-KL, HPR1-HA::KanMX4 | This study |
| *npl3* shuffle *MEX67-HA* | MATa, ura3-1, trp1-1, his3-11,15, leu2-3,112, ade2-1, can1-100 GAL+, NPL3::HIS3, pRS316-NPL3, MEX67-HA::KanMX4 | This study |
| *npl3* shuffle *HPR1-HA* | MATa, ura3-1, trp1-1, his3-11,15, leu2-3,112, ade2-1, can1-100 GAL+, NPL3::HIS3, pRS316-NPL3, HPR1-HA::KanMX4 | This study |
| *npl3* shuffle *SUB2-FTpA* | MATa, ura3-1, trp1-1, his3-11,15, leu2-3,112, ade2-1, can1-100 GAL+, NPL3::HIS3, pRS316-NPL3 | This study |
| *npl3* shuffle THO1-FTpA | MATa, ura3-1, trp1-1, his3-11,15, leu2-3,112, ade2-1, can1-100 GAL+, NPL3::HIS3, pRS316-NPL3 | This study |
| *npl3* shuffle *MEX67-FTpA* | MATa, ura3-1, trp1-1, his3-11,15, leu2-3,112, ade2-1, can1-100 GAL+, NPL3::HIS3, pRS316-NPL3 | This study |
| *npl3* shuffle *HPR1-FTpA* | MATa, ura3-1, trp1-1, his3-11,15, leu2-3,112, ade2-1, can1-100 GAL+, NPL3::HIS3, pRS316-NPL3 | This study |

### Supplementary Table S10. Plasmids

| Plasmid | Description | Reference |
|---|---|---|
| pBS1479 | For genomic C-terminal TAP-tagging (CBP-TEV-2x protein A), *TRP1-KL* | Puig O. 2001 |
| pFA6a-FTpA-HIS3MX4 | For genomic C-terminal FTpA-tagging (FLAG-TEV-2x protein A), *HIS3MX4* | Kressler D. 2012 |
| pRS316-*NPL3* | pBlueScript based yeast centromere vector with *URA3* marker and ORF + 500 bp of 5’ and 300 bp of 3’ UTR of *NPL3* | This study |
| pRS315 | pBlueScript based yeast centromere vector with *LEU2* marker | Sikorski R.S. 1989 |
| pRS315-*NPL3* | ORF + 500 bp of 5’ and 300 bp of 3’ UTR of *NPL3* was cloned into pRS315 | This study |
| pRS315-*npl3-F160Y,F162Y,F165Y* | *npl3-F160Y,F162Y,F165Y* | This study |
| pRS315-*npl3-F160Y,F162Y* | *npl3-F160Y,F162Y* | This study |
| pRS315-*npl3-F160Y,F165Y* | *npl3-F160Y,F165Y* | This study |
| pRS315-*npl3-F162Y,F165Y* | *npl3-F162Y,F165Y* | This study |
| pRS315-*npl3- F160Y* | *npl3-F160Y* | This study |
| pRS315-*npl3-F162Y (RRM1)* | *npl3-F162Y (RRM1)* | This study |
| pRS315-*npl3- F165Y* | *npl3-F165Y* | This study |
| pRS315-*npl3-P196D,A197D (Linker)* | *npl3-P196D,A197D (Linker)* | This study |
| pRS315-*npl3-F245I (RRM2)* | *npl3-F245I (RRM2)* | This study |
| pRS315-*npl3-F245Y* | *npl3-F245Y* | This study |

| | | |
|---|---|---|
| pRS315-*npl3-RRM1-Linker* | *npl3-F162Y,P196D,A197D* | This study |
| pRS315-*npl3-RRM1-RRM2* | *npl3-F162Y,F245I* | This study |
| pRS315-*npl3-Linker-RRM2* | *npl3-P196D,A197D,F245I* | This study |
| pRS315-*npl3-RRM1-Linker-RRM2* | *npl3-F162Y,P196D,A197D,F245I* | This study |
| pRS315-*TAP-NPL3* | *2x prot A-TEV-CBP-NPL3* | This study |
| pRS315-*TAP-npl3-RRM1* | *2x prot A-TEV-CBP-npl3-F162Y* | This study |
| pRS315-*TAP-npl3-Linker* | *2x prot A-TEV-CBP-npl3-P196D,A197D* | This study |
| pRS315-*TAP-npl3-F245I (RRM2)* | *2x prot A-TEV-CBP*-npl3*-F245I* | This study |
| pRS315-*TAP-npl3-F245Y* | *2x prot A-TEV-CBP*-npl3*-F245Y* | This study |
| pRS315-*TAP-npl3-RRM1-Linker* | *2x prot A-TEV-CBP-npl3-F162Y,P196D,A197D* | This study |
| pRS315-TAP-*npl3-RRM1-RRM2* | *2x prot A-TEV-CBP-npl3-F162Y,F245I* | This study |
| pRS315-TAP-*npl3-Linker-RRM2* | *2x prot A-TEV-CBP-npl3-P196D,A197D,F245I* | This study |
| pRS315-TAP-*npl3-RRM1-Linker-RRM2* | *2x prot A-TEV-CBP-npl3-F162Y,P196D,A197D,F245I* | This study |
| pRS425 | *high copy number plasmid, 2μ, with LEU2 maker* | (5) |
| pRS425-*NPL3* | *ORF + 500 bp of 5' and 300 bp of 3' UTR of NPL3 was cloned into pRS425* | This study |
| pRS425-*npl3-RRM1* | *npl3-F162Y (RRM1)* | This study |
| pRS425-*npl3-Linker* | *npl3-P196D,A197D (Linker)* | This study |
| pRS425-*npl3-RRM2* | *npl3-F245I (RRM2)* | This study |
| pYM14 | For genomic C-terminal HA-tagging, KanMX4 | Janke C., 2004 |
| pT7-His6-TEV-NPL3 (RRMs, 121-280) | For recombinant expression of both RRM domains of Npl3; His6-TEV-Npl3 (aa121-aa280) | This study |
| pT7-His6-TEV-*npl3-RRM1* (RRMs, 121-280) | For recombinant expression of both RRM domains of npl3-RRM1 | This study |
| pT7-His6-TEV-*npl3-Linker* (RRMs, 121-280) | For recombinant expression of both RRM domains of npl3-Linker | This study |
| pT7-His6-TEV-*npl3-RRM2* (RRMs, 121-280) | For recombinant expression of both RRM domains of npl3-RRM2 | This study |
| pT7-His6-TEV-NPL3 (RRMs, 120-280) | Recombinant expression and purification of Npl3-RRM1,2 for NMR experiments | This study |
| pT7-His6-TEV-NPL3-RRM1 only | Recombinant expression and purification of Npl3-RRM1 for NMR titration experiments | This study |
| pT7-His6-TEV-NPL3-RRM2 only | Recombinant expression and purification of Npl3-RRM2 for NMR titration experiments | This study |
| npl3-D135C & C211S | Single cysteine mutant D135C of Npl3-RRM1,2 for PRE experiments | This study |
| npl3-E176C & C211S | Single cysteine mutant E176C of Npl3-RRM1,2 for PRE experiments | This study |
| npl3-N185C & C211S | Single cysteine mutant N185C of Npl3-RRM1,2 for PRE experiments | This study |

| | | |
|---|---|---|
| npl3-D236C & C211S | Single cysteine mutant D236C of Npl3-RRM1,2 for PRE experiments | This study |

### Supplementary Table S11. Primers

| **Primers used for genomic tagging** | **Sequence (5' to 3')** |
|---|---|
| HPR1-FTpA fwd | GCAGCTACTTCGAACATTTCTAATGGTTCATCTACCCAAGATATGAAAGATCA CGACGGTGACTAC |
| HPR1-FTpA rev | TGCATGAATTTCTTATCAGTTTAAAATTTCTATTAAGAGGATAATTTAATCGAT GAATTCGAGCTCG |
| THO1-FTpA fwd | TCCAGAGTAAGTAAAAACAGGAGAGGCAACCGCTCTGGTTACAGAAGAGATC ACGACGGTGACTAC |
| THO1-FTpA rev | GAACCGAAACTAGAATGAAAAACTCCACCAAAACGGCTTGAGCCTTTAATCG ATGAATTCGAGCTCG |
| NAB2-FTpA fwd | CCTCCGCAAACCAGTTTTACGCACCAAGAACAAGATACGGAAATGAACGATC ACGACGGTGACTAC |
| NAB2-FTpA rev | AAGGGTCACAGGAACATGAATTTCGTTCCGTGATTTTAATAGTAATCAATCGA TGAATTCGAGCTCG |
| MTR2-FTpA fwd | TTTAACTATAACATGGTTTACAAACCCGAGGATTCTCTGCTAAAAATTGATCA CGACGGTGACTAC |
| MTR2-FTpA rev | TTTCTATATAATTTTGTTTTTCCCTGGTAGAGCGGAATCTTCCCACTAATCGAT GAATTCGAGCTCG |
| CBC2-FTpA fwd | AGACCAGGTTTCGATGAAGAAAGAGAAGATGATAACTACGTACCTCAGGATC ACGACGGTGACTAC |
| CBC2-FTpA rev | ATATATCTGTGTGTAGAATCTTTCTCAGATATAAATTGATTGATTCTAATCGAT GAATTCGAGCTCG |
| THP1-FTpA fwd | AATGAACGAATCACCAAGATGTTTCCTGCCCATTCTCACGTTCTTTGGGATCA CGACGGTGACTAC |
| THP1-FTpA rev | TGATATGTAGATATATATAGGAAATAGAAGAGAAGGAGCGACATTATGATCGA TGAA TTCGAGCTCG |
| CBC2-TAP fw | AGACCAGGTTTCGATGAAGAAAGAGAAGATGATAACTACGTACCTCAGTCCA TGGAAAAGAGAAG |
| CBC2-TAP rev | TATATATATCTGTGTGTAGAATCTTTCTCAGATATAAATTGATTGATTCTATAC GACTCACTATAGGG |
| MEX67-HA fwd | TAAACTGTATATTTTTTGTGATACTGTGCGGCTGAAACAGGGAACAATATCAT TAATCGATGAATTCGAGCTCG |
| MEX67-HA rev | AAAGGGTTTTCAGAGTAGCATGAATGGCATCCCTAGAGAAGCATTTGTGCAG TTCCGTACGCTGCAGGTCGAC |
| HPR1-HA fwd | GCTACTTCGAACATTTCTAATGGTTCATCTACCCAAGATATGAAACGTACGCT GCAGGTCGAC |
| HPR1-HA rev | ATGAATTTCTTATCAGTTTAAAATTTCTATTAAGAGGATAATTTAATCGATGAA TTCGAGCTCG |
| SUB2-FTpA fwd | GCTGAATTCCCAGAAGAAGGCATTGATCCGTCCACTTATTTGAATAATGATC ACGACGGTGACTAC |
| SUB2-FTpA rev | AAAATCTTTATATAATCTATATAAAAACGTATCTTTTTTCCTTTATTAATCGATG AATTCGAGCTCG |
| MEX67-FTpA fwd | TTTCAGAGTAGCATGAATGGCATCCCTAGAGAAGCATTTGTGCAGTTCGATCA CGACGGTGACTAC |
| MEX67-FTpA rev | TATATTTTTTGTGATACTGTGCGGCTGAAACAGGGAACAATATCATTAATCGA TGAATTCGAGCTCG |
